# Human engineered skeletal muscle of hypaxial origin from pluripotent stem cells with advanced function and regenerative capacity

**DOI:** 10.1101/2021.07.12.452030

**Authors:** Mina Shahriyari, Md Rezaul Islam, M. Sadman Sakib, Anastasia Rika, Dennis Krüger, Lalit Kaurani, Harithaa Anandakumar, Orr Shomroni, Matthias Schmidt, Gabriela Salinas, Andreas Unger, Wolfgang A. Linke, Jana Zschüntzsch, Jens Schmidt, André Fischer, Wolfram-Hubertus Zimmermann, Malte Tiburcy

**Affiliations:** Institute of Pharmacology and Toxicology, University Medical Center Göttingen, Göttingen, Germany.; DZHK (German Center for Cardiovascular Research), partner site Göttingen, Germany; Department for Epigenetics and Systems Medicine in Neurodegenerative Diseases, German Center for Neurodegenerative Diseases (DZNE) Göttingen, Göttingen, Germany; NGS Integrative Genomics Core Unit, Institute of Human Genetics, University Medical Center Göttingen, Göttingen, Germany; Department of Neurology, University Medical Center Göttingen, Göttingen, Germany; Institute of Physiology II, University of Münster, D-48149 Münster, Germany; Department of Neurology and Pain Treatment, University Hospital of the Medical School Brandenburg, Immanuel Klinik Rüdersdorf, Rüdersdorf bei Berlin, Germany; Cluster of Excellence "Multiscale Bioimaging: from Molecular Machines to Networks of Excitable Cells" (MBExC), University of Göttingen, Germany; Fraunhofer Institute for Translational Medicine and Pharmacology (ITMP), Göttingen, Germany

**Keywords:** Skeletal muscle organoid, tissue engineering, limb muscle, hypaxial dermomyotome, satellite cells, regeneration, spinal neurons, motor end plate

## Abstract

Human pluripotent stem cell derived muscle models show great potential for translational research. Here, we describe developmentally inspired methods for derivation of skeletal muscle cells and their utility in three-dimensional skeletal muscle organoid formation as well as skeletal muscle tissue engineering. Key steps include the directed differentiation of human pluripotent stem cells to embryonic muscle progenitors of hypaxial origin followed by primary and secondary fetal myogenesis into hypaxial muscle with development of a satellite cell pool and evidence for innervation *in vitro*. Skeletal muscle organoids faithfully recapitulate all steps of embryonic myogenesis in 3D Tissue engineered muscle exhibits organotypic maturation and function, advanced by thyroid hormone. Regenerative competence was demonstrated in a cardiotoxin injury model with evidence of satellite cell activation as underlying mechanism. Collectively, we introduce a hypaxial muscle model with canonical properties of *bona fide* skeletal muscle *in vivo* to study muscle development, maturation, disease, and repair.

## Introduction

Pluripotent stem cell (PSC)-derived organotypic cultures with structural and functional properties of native human tissue are increasingly utilized for disease modeling and drug screening applications (Tachibana, 2018). Organotypic skeletal muscle cultures are highly sought after, because of the central role of skeletal muscle in disease (e.g., myopathies) and drug effects (e.g., insulin). Early studies have demonstrated that it only requires MyoD overexpression in fibroblasts cells to recreate skeletal muscle cells (Davis et al., 1987). Alternatively, muscle stem cells can be isolated from muscle biopsies, but rapidly lose their stem cell properties with expansion often requiring immortalization to provide sufficient numbers of consistent cell quality (Mamchaoui et al., 2011; Striedinger et al., 2021). In those muscle models developmental information on muscle origin is not existent or lost. Deriving skeletal muscle cells from human PSC via directed differentiation closely recapitulating embryonic myogenesis may overcome these shortcomings.

The derivation of skeletal muscle cells from PSC has been demonstrated previously either by transfection or transduction of myogenic transgenes (Albini et al., 2013; Darabi et al., 2012; Goudenege et al., 2012; Kim et al., 2017; Rao et al., 2018; Tedesco et al., 2012; Young et al., 2016) or directed, transgene-free differentiation under controlled growth factors or small molecules stimulation (Borchin et al., 2013; Caron et al., 2016; Chal et al., 2016; Chal et al., 2015; Choi et al., 2016; Shelton et al., 2016; Xi et al., 2017). Recently, more advanced skeletal muscle organoids have been introduced that recapitulate characteristic steps of embryonic neuromuscular co-development (Faustino Martins et al., 2020; Mazaleyrat et al., 2020).

The embryonic development of skeletal muscle is a complex process with intricate interplay of decisive transcriptional programs (Buckingham, 2017). Following the specification of presomitic (paraxial) mesoderm from epiblast/neuromesodermal progenitors (NMPs), trunk and limb muscle derives from developing somites. The myogenic structure to form first in the developing somite is the dermomyotome, which can be anatomically divided into dorsomedial (epaxial) and ventrolateral (hypaxial) compartments giving rise to skeletal muscle of back and trunk/limb, respectively. Hypaxial PAX3+ dermomyotomal progenitors cells, characterized by LBX1 and SIM1 expression, migrate into the limb bud to form limb muscle (Buckingham and Mayeuf, 2012; Coumailleau and Duprez, 2009). Further myogenic differentiation is then proceeded with embryonic primary and fetal secondary myofiber formation (Biressi et al., 2007). Recent work has contributed significantly in dissecting transcriptome profiles and cell composition in developing human limb muscle (Xi et al., 2017; Xi et al., 2020).

Several studies have applied tissue engineering methods to generate skeletal muscle from human pluripotent stem cell-derived cells *in vitro* (Maffioletti et al., 2018; Rao et al., 2018; Xu et al., 2019) collectively suggesting a potential of 3D skeletal muscle for disease modeling and regenerative medicine. However, the functional output of *in vitro* muscle is still far from postnatal muscle even though improvements have been demonstrated using a specific cell culture supplement (Xu et al., 2019). For the only transgene-free model published so far, muscle function was not reported (Maffioletti et al., 2018). In a previous study using rat primary myocytes, our group demonstrated that the application of collagen/Matrigel^®^ hydrogels in combination with isometric loading generates Engineered Skeletal Muscle (ESM) with physiological function and a regenerative satellite cell niche *in vitro* (Tiburcy et al, 2019).

Here, we report a transgene-free and completely serum-free human muscle protocol that closely follows developmental, tissue-specific paradigms to derive hypaxial muscle of limb and trunk which shows regenerative as well as innervation capacity. Human ESM respond to developmentally relevant cues, such as triiodothyronine, with an advanced maturation, demonstrating physiological growth potential and establishing the groundwork for further optimization of *in vitro* engineered human skeletal muscle for applications in developmental studies, disease modelling, and muscle regeneration research.

## Results

### Directed differentiation of hypaxial skeletal myocytes from human pluripotent stem cells

To robustly generate human skeletal muscle, we followed several developmental paradigms, which were first optimized in monolayer cultures by directed skeletal muscle differentiation (final optimized protocol in **Figure 1A**). We reasoned that a modification of BMP (inhibition) and Wnt (activation) signaling pathways, previously identified as crucial for myogenesis (Chal et al., 2016; Shelton et al., 2016; Shelton et al., 2014), would be a good starting point. Whereas Wnt activation by CHIR99021 (10 μmol/L; GSK-3α/β inhibition with an IC50 ≤10 nmol/L) serves as a more generic mesoderm induction measure (Lian et al., 2012; Mendjan et al., 2014; Tiburcy et al., 2017), it is the parallel inhibition of BMP with LDN 193189 (0.5 μmol/L; ALK2 an ALK3 inhibition with an IC50 ≤30 nmol/L), which is instrumental in specifying *MSGN1*+ paraxial mesoderm (Miura et al., 2006) versus *MESP1*+ lateral plate mesoderm (**Figure 1B**). Following paraxial mesoderm induction, Notch 1 signaling was blocked with DAPT (10 µmol/L; γ-secretase inhibition;) in the presence of FGF2 and HGF to support somitogenesis and inhibit Notch 1 dependent epaxial dermomyotomal progenitors (Bladt et al., 1995; Mayeuf-Louchart et al., 2014; Rios et al., 2011). The resulting formation of hypaxial progenitors, which give rise to limb and trunk muscle, was confirmed by the expression of *PAX3*, *SIM1, LBX1,* and lack of *EN1* (**Figure 1C**). Considering that continuous Notch inhibition prevents differentiation of *PAX3*+ cells to early myoblasts, DAPT was then discontinued from day 13 of differentiation to allow for the expression of myogenic regulatory factors (MRFs) such as *PAX7*, *MYOD1* and *MYOG* as well as sarcomeric transcripts such as *ACTN2* (**Figure 1D**; (Choi et al., 2016; Hirsinger et al., 2001)). Immunostaining confirmed the specific stages of skeletal muscle differentiation with timed expression of characteristic myogenic regulatory factors (PAX3, PAX7, MYOD1, MYOG) on protein level. In addition, the sarcomeric actinin (ACTN2) staining indicated highly efficient skeletal myocyte generation with this protocol (**Figure 1E**).

**Figure 1.**
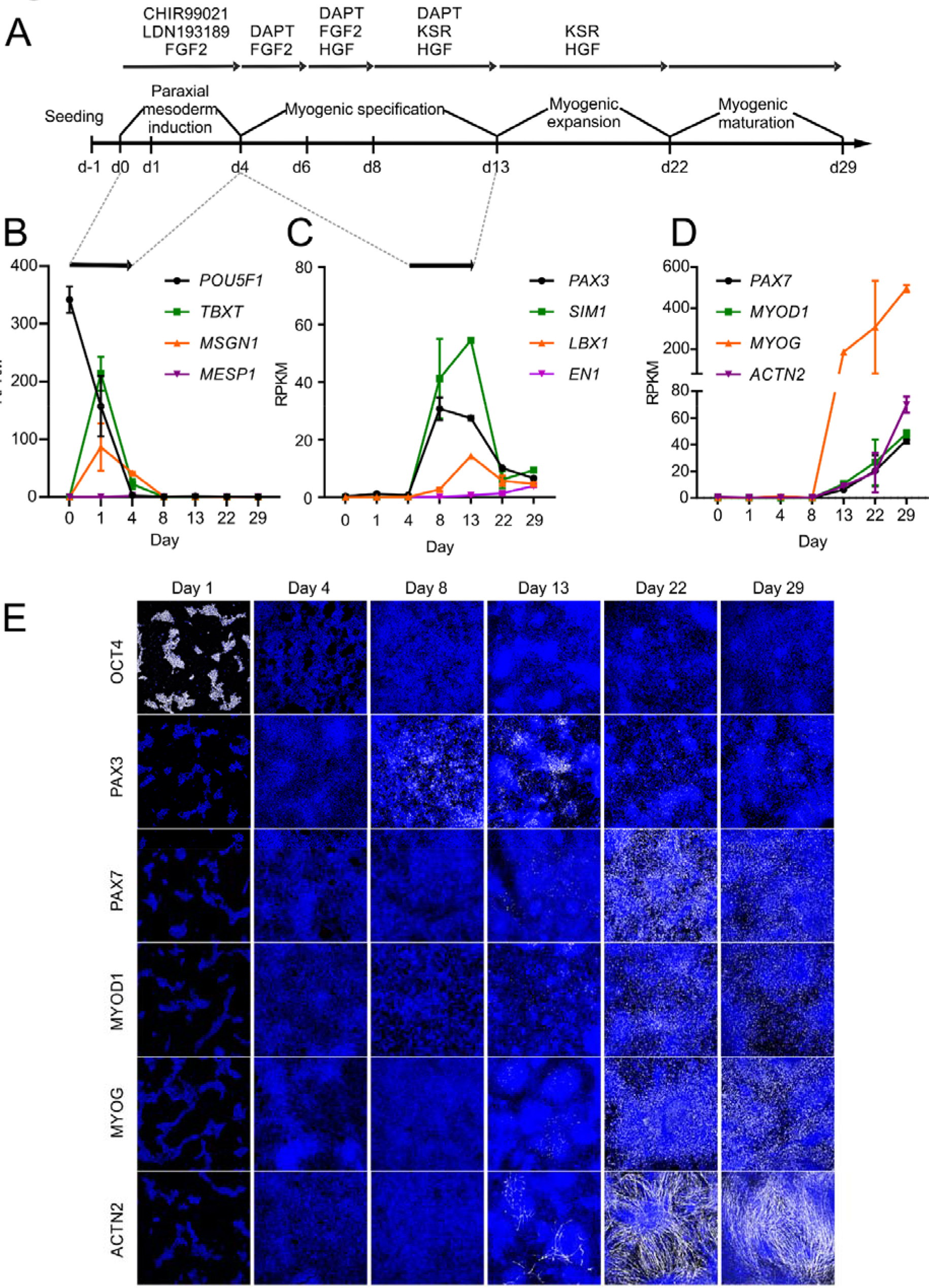
Hypaxial skeletal myocyte differentiation from human pluripotent stem cells. **(A)** Summary of the protocol for directed skeletal muscle differentiation from pluripotent stem cells (PSC) indicating the sequence and the timing of factor addition to modulate specific signaling pathways involved in skeletal myogenesis. Reads Per Kilobase Million (RPKM) of signature genes for **(B)** pluripotency (*POU5F1*), naïve mesoderm (TBXT), paraxial mesoderm (*MSGN1*), and lateral plate mesoderm (*MESP1*); **(C)** dermomyotome formation (*PAX3*), hypaxial (*SIM1, LBX1*) and epaxial (*EN1*) dermomyotome, and **(D)**, myogenic regulatory factors (*PAX7*, *MYOD1* and *MYOG*) and structural assembly (*ACTN2*), during skeletal muscle differentiation from human PSCs; n = 2-4/time point. (**E)** Immunostaining of OCT4, PAX3, PAX7, MYOD1, MYOGENIN, sarcomeric ACTININ (in gray), and nuclei (blue) at indicated time points of skeletal muscle differentiation. Scale bar: 500 µm.

### Myocyte differentiation closely follows embryonic skeletal muscle development *in vivo*

To obtain more insight into the global developmental patterns of skeletal muscle differentiation *in vitro* we subjected the RNAseq data obtained at selected time points (**Figure 2A**) to bioinformatic analyses. Unbiased clustering clearly separated the distinct time points into mesoderm induction (days 0, 1 and 4), myogenic specification (days 8 and 13), early (days 22 and 29) and advanced (day 60) myogenic maturation (**Figure 2B**). Further clustering the genes by weighted co-expression analysis identified 22 gene clusters with remarkable overlap to the biological processes of skeletal muscle differentiation characterized by the expression of developmental signature genes (**Figure 2C,D**, **Supplementary Table 1**). Coinciding with the loss of pluripotency, we observed an increase in primitive streak transcripts (*MIXL*). Expression of *TBX6* (cluster “brown”) and the segmentation gene *HES7* (cluster “red”) indicated robust paraxial mesoderm formation and patterning, which was followed by dermomyotomal progenitor transcript *FOXC2* (cluster “turquoise”; **Figure 2D**). By day 8 of differentiation robust *PAX3, TWIST1* and *SIM1* expression indicated the formation of the hypaxial dermomyotomal cells (cluster “salmon”) and migrating limb progenitors (*SIX1, MET, MEOX2,* cluster “black”). This was followed by an increase in genes indicative of myoblast generation and fusion (*MYMX, NGFR, ERBB3*) and an increase of more mature transcripts such as *MYF6, TTN, MYL3* (cluster “blue”).

**Figure 2.**
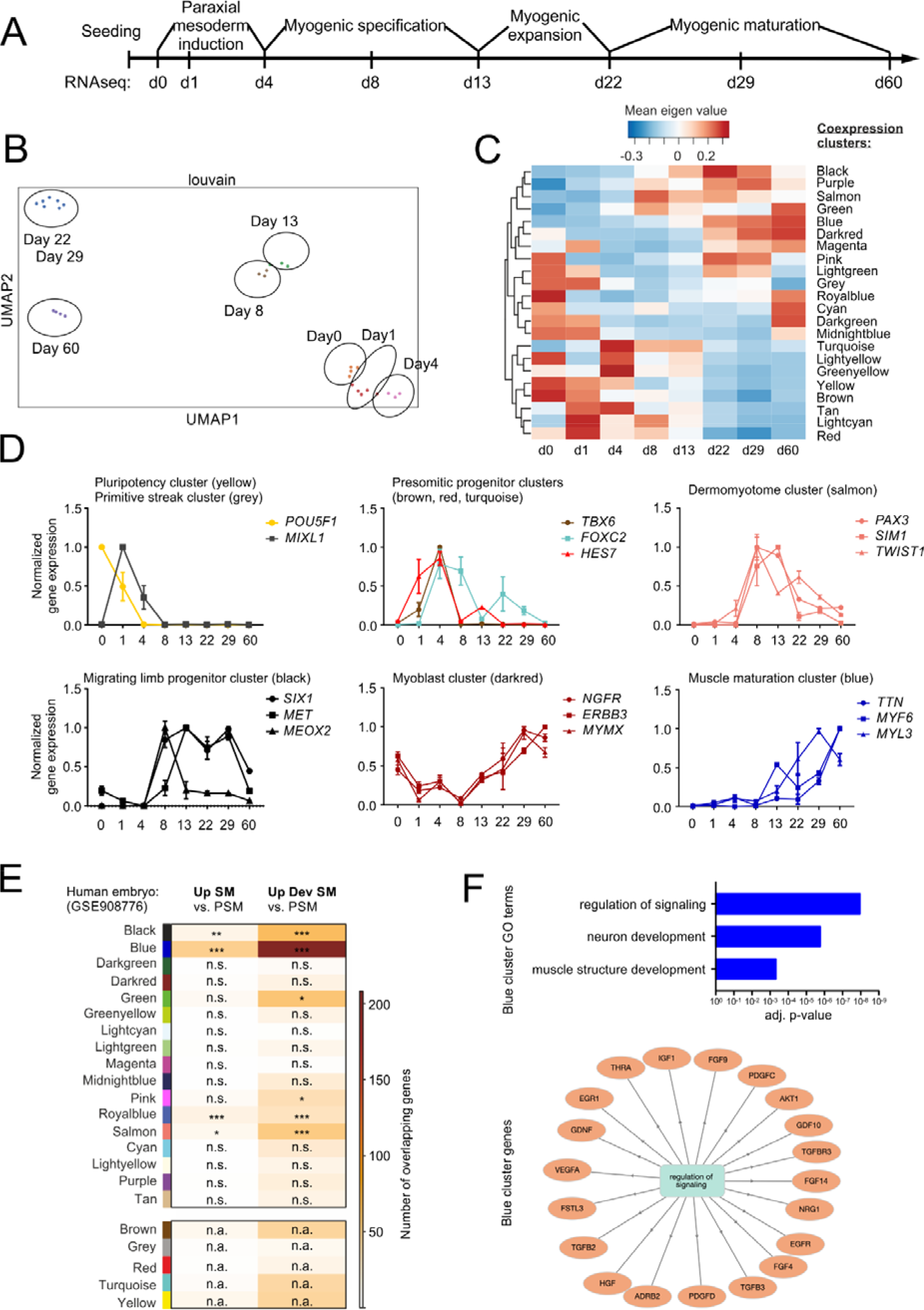
Developmental transcriptome patterns in PSC skeletal myocyte differentiation. **(A)** Scheme of skeletal muscle differentiation from hPSCs with sampling time points for RNA sequencing. **(B)** Unsupervised clustering of the samples from different time points. **(C)** Weighted coexpression analysis identified 22 cluster of genes with similar expression dynamics (coexpression clusters); a heatmap of mean eigen values is displayed. Clusters are generically labeled by colors. **(D)** Normalized expression levels (RPKM) of indicated signature genes in identified coexpression clusters, n = 2-4/time point. **(E)** Developmentally regulated genes were identified based on a published human embryonic muscle data set (Xi et al. 2017). The table indicates the overlap of coexpression cluster genes to genes regulated between presomitic mesoderm (PSM) and nascent somite (SM) or presomitic mesoderm (PSM) and developed somite (Dev SM). Overlap is graded as either not significant (n.s.), p<0.05 (*), p<0.01 (**), or p<0.001 (***) by Fishers’s exact test. The color codes for the number of genes overlapping. Early developmental clusters that cannot be represented in the embryo data set are labelled as not applicable (n.a.). **(F)** GO terms specifically enriched in coexpression cluster blue **(top panel)**. List of genes associated with “regulation of signaling” in coexpression cluster blue **(bottom panel)**.

We next asked if the identified gene clusters overlap with developmentally regulated genes of human embryonic muscle. We made use of a published data set (GSE908776) containing the transcriptomes of embryonic presomitic mesoderm (PSM), nascent somite (SM), and developed somite (SM dev) from human embryonic tissue (Xi et al., 2017). Interestingly, several of the bioinformatically identified clusters from skeletal myogenesis *in vitro* showed significant overlap with the data obtained from embryonic development *in vivo* (**Figure 2E**). Note that the very early developmental gene clusters (i.e., paraxial mesoderm stage and earlier) are not represented in the embryo data set and therefore cannot overlap (labelled as not applicable, n.a.).

We then utilized the co-expression analysis to dissect processes coinciding with particular stages of muscle development. We focused on the “blue” cluster as this showed the highest overlap with the developed somite *in vivo* (**Figure 2E**) and may therefore contain transcripts that support muscle differentiation and maturation. Interestingly, cluster “blue” was highly enriched in signaling transcripts (**Figure 2F**). Among them we identified several signaling pathways that have been associated with muscle maturation such as NRG1 (Selvaraj et al., 2019), IGF1/VEGF (Xu et al., 2019), and thyroid hormone (Larsson et al., 1994; Schiaffino et al., 1988; Simonides and van Hardeveld, 2008) indicating that our protocol emulates central mechanisms of muscle development *in vivo*.

### Myocyte and non-myocyte populations of embryonic myogenesis *in vitro*

As skeletal muscle differentiation from PSC may yield heterogenous cell populations (Xi et al., 2020) we aimed to characterize the cell composition on single cell level. *Bona-fide* skeletal myogenic markers were quantified by immunostaining (**Figure 3A**). Differentiated cultures at day 22 contained a myogenic cell population consisting of 43±4% PAX7+, 52±2% MYOD1+, and 49±4% MYOGENIN+ (n=9-13 differentiations) cells (**Figure 3B**). Per input PSC we obtained 61±5 (n=10) differentiated skeletal myogenic cells. Flow cytometry confirmed the quantification of immunostaining data and showed comparable efficiency for 1 human embryonic stem cell (HESC) and 4 different induced pluripotent stem cell (iPSC) lines supporting the robustness and reproducibility of the protocol (**Supplementary Figure 1**). We then investigated day 22 cultures by single nuclei sequencing to gain further insight into the cell composition. 8 cell populations were separated by unsupervised clustering (**Figure 3C**). To identify myogenic cells a panel of genes (muscle genes, **Supplementary Table 2**) was extracted. This panel identified 3 myogenic cell populations accounting for 45% of the total cell population (**Figure 3D, E**). Non-muscle cells were identified as mesenchymal progenitor cells (41%), neuroectodermal progenitor cells (9%), and neurons (5%; **Figure 3E**). We did not identify a specific Schwann cell (*CHD19+*), (pre-)chondrocyte (*SHOX2+*), or tenogenic cell (*TNMD+)* cell population in contrast to other protocols (Xi et al., 2020). The myogenic cells mainly consisted of *PAX7+* progenitors and relatively few matured *MYH3+* myoblasts in line with the early time point of analysis. The neuronal population transcript signatures suggested the presence of bona fide neurons (*MAPT+*) and neuroectodermal progenitor cells (*SOX2+/PAX3+/PAX7+*). The mesenchymal cells showed an expression pattern consistent with limb fibro-adipogenic progenitor (FAP) cells with combined expression of *PDGFRA, MEOX2* and brown fat transcription factor *EBF2* (Rajakumari et al., 2013; Reijntjes et al., 2007; Uezumi et al., 2010; Xi et al., 2020). These data indicate that the differentiation protocol not only recapitulates the embryonic development of muscle cells but that of essential non-muscle cells.

**Figure 3.**
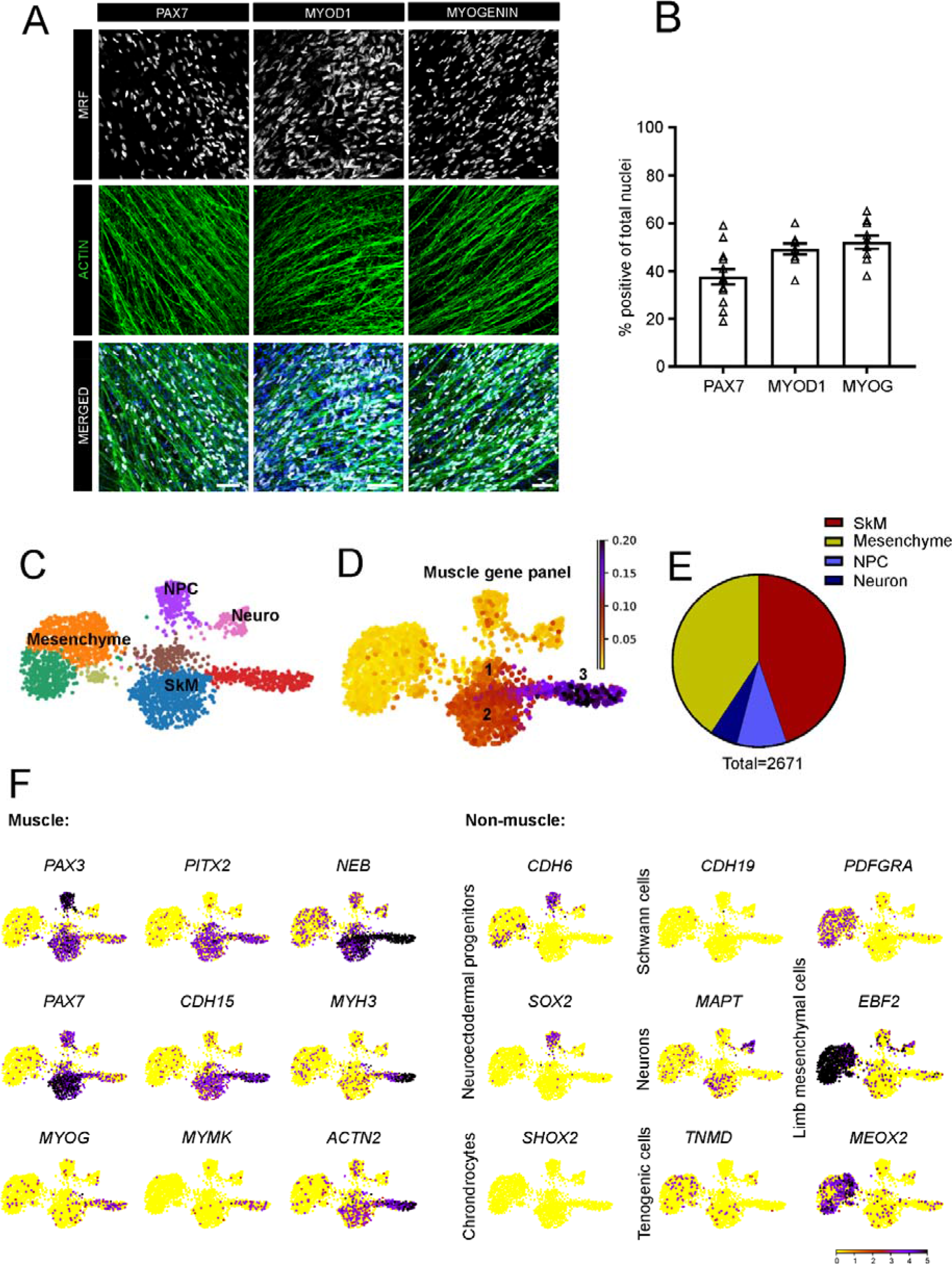
Cellular composition of differentiated skeletal myogenic cultures. **(A)** Representative immunostaining of myogenic regulatory factors (MRF): PAX7, MYOD1 and MYOGENIN (gray), f-ACTIN (green) and nuclei (blue) in 22 days old skeletal muscle cultures from TC1133 (iPSC 1) line; Scale bars: 50 µm. **(B)** Quantification of nuclei positive for PAX7, MYOD1 and MYOGENIN in 22 days old myogenic cultures from HES2 and from TC1133 (iPSC 1) lines; n = 9 -13 differentiations. **(C)** Unsupervised clustering (UMAP) of single nuclei transcriptomes from a day 22 skeletal muscle culture. **(D)** A muscle gene panel identifies 3 myogenic cell clusters (skeletal muscle cells, SkM). **(E)** Quantification of skeletal muscle cells (SkM), neuroectodermal progenitor cells (NPC), neurons, and mesenchymal progenitor cells (limb mesenchyme) of a total of 2,671 nuclei analyzed. **(F)** Expression levels of representative muscle-related genes and non-muscle genes.

### Directing iPSC to force-generating skeletal muscle organoids

As 2D cultures of skeletal myocytes do not develop the spatial and structural organization of skeletal muscle *in vivo* (Afshar Bakooshli et al., 2019) we next asked if the muscle generation process could be fully recapitulated in a 3D environment. To test this, we embedded undifferentiated iPSC in a collagen/Matrigel hydrogel [adapted from our previous skeletal muscle engineering work in rodents (Tiburcy et al. 2019)] and subjected them to the for the human PSC established differentiation protocol (**Figure 1A**) to obtain skeletal muscle organoids (SMOs; **Figure 4A**). 3D SMO development was supported by the casting of the iPSC/matrix mixture in circular molds. SMO rings formed within 24 h after casting, Transfer of SMO onto holders of defined distance on culture day 22 supported further maturation under defined mechanical strain (**Figure 4A**). We next asked if the temporal sequence of muscle cell development was comparable to 2D differentiation. RNA expression analyses showed a decrease in pluripotency genes (*POU5F1*), increase of paraxial mesoderm (*MSGN1*), dermomyotome (*PAX3*), and muscle progenitors expressing *PAX7, MYOD1*, and *MYOG* similar as in 2D cultures (**Figure 4B**). Importantly, we observed an identical pattern of hypaxial progenitor specification expressing *SIX1* and *SIM1* with only low levels of *EN1.* In addition, robust increases in *NFIX* and *ENO3* expression suggested secondary myogenesis in maturing SMOs between day 22 and 52 (**Figure 4C**). Compatible with the expression data significant amounts of differentiated, multinucleated muscle fibers were identified by immunostaining. F-actin and dystrophin-associated glycoprotein, ’Q-dystroglycan staining on SMO cross sections indicated that 48±6% (n=3) of the total cross sectional area was populated with muscle cells (**Figure 4D**). Muscle cells were aligned and cross striated indicating a well-developed sarcomeric structure (**Figure 4E**). In line with the transcriptome data of single cell nuclei in monolayer cultures, SMOs also contained neurofilament positive cells co-labeled for the motoneuron marker SMI32 (**Figure 4E**), suggesting innervation in line with recently reported data (Faustino Martins et al., 2020). Maturing SMOs demonstrated spontaneous contractions (**Video 1**) and robust force generation at day 50. At 1 Hz electrical stimulation SMOs generated single twitches whereas at higher frequencies tetanic contractions were observed which were maximal at 100 Hz (1.1±0.1 mN, n=9; **Figure 4F,G**). In summary, we demonstrate the generation of functional muscle of hypaxial origin directly from human PSC in a novel 3D organoid approach.

**Figure 4.**
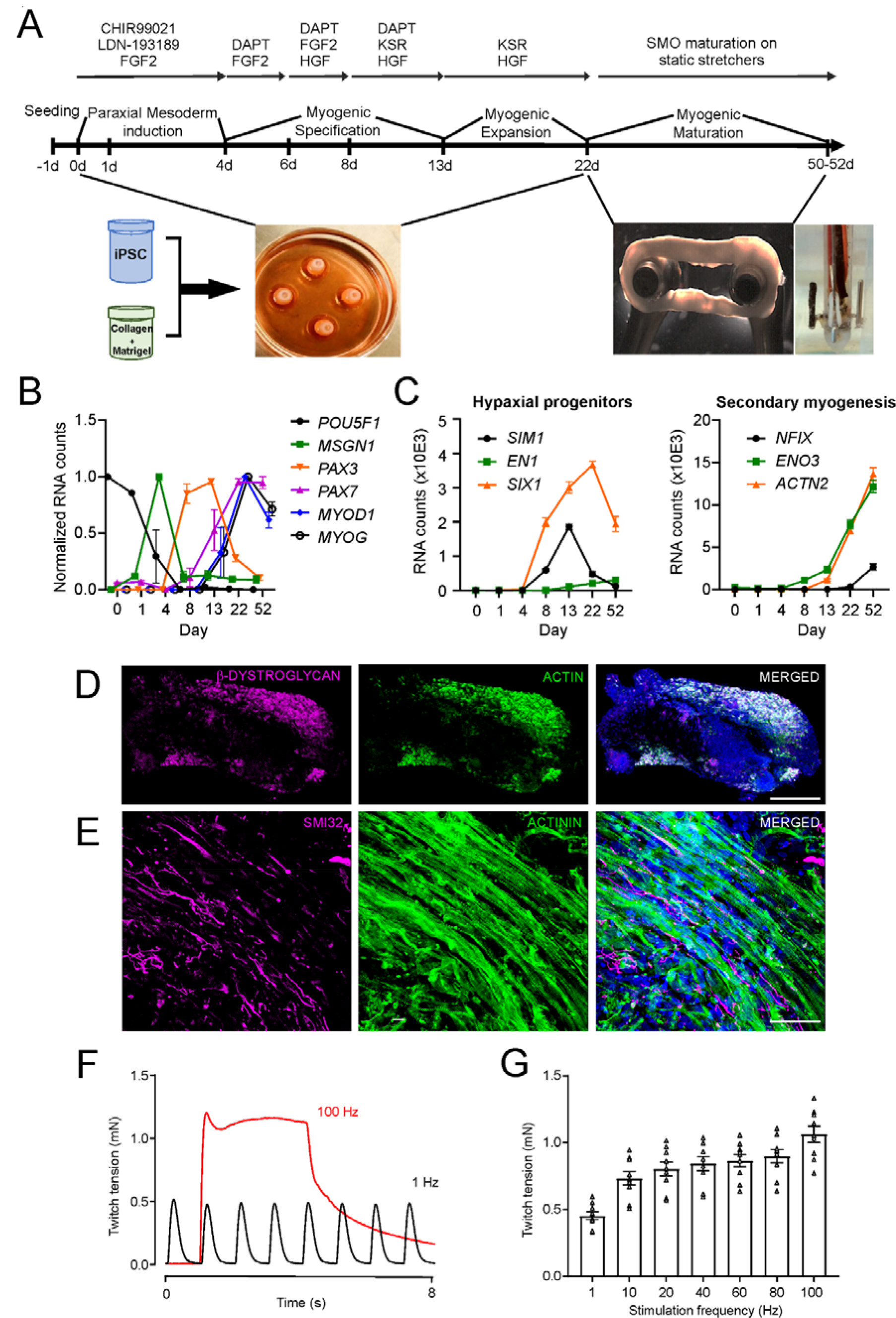
Generation of functional hypaxial skeletal muscle organoids. **(A)** Skeletal muscle organoids (SMO) were generated from iPSC mixed with collagen type 1 and Matrigel™ in a ring-shaped hydrogel. After consolidation in PDMS casting molds, SMOs were directed towards skeletal muscle using the protocol established in monolayer cultures. Following functional maturation under isometric load for 3-5 wks (day 22 to 52), twitch tension (TT) was measured under isometric conditions in a thermostatted organ bath. Scale bar: 5 mm. **(B)** RNA expression of indicated genes (RNA counts measured by nCounter) normalized to minimal and maximal expression at different days of SMO culture. **(C)** RNA expression (RNA counts measured by nCounter) of marker genes of hypaxial progenitor cells (left panel) and secondary myogenesis (right panel). **(D)** Immunostaining of the total muscle area in cross sections of BSM with β*-*DYSTROGLYCAN (magenta), ACTIN+ (green), and Nuclei (blue). Scale bar: 500 µm. **(E)** Immunostaining of neurofilament heavy SMI32 (magenta), sarcomeric ɑ: -ACTININ (green) and nuclei (blue) in longitudinal sections of BSM. Scale bar: 50 µm. **(F)** Representative force traces of 4 weeks old SMO with electrical stimulation at 1 Hz (black curve) and at 100 Hz (red curve). **(G)** Quantification of the twitch tension (TT) generated by 4 weeks old SMO in response to increasing stimulation frequencies; n = 9/group.

### Engineering human muscle with advanced muscle function

As an alternative and more controlled tissue engineered approach to generate human skeletal muscle, we tested whether already differentiated PSC-derived skeletal myocyte populations obtained from the 2D directed differentiation can be assembled into engineered skeletal muscle (PSC ESM), following a protocol developed by our group for heart muscle engineering (Tiburcy et al. 2017) and more recently adapted to skeletal muscle engineering in a rodent model (Tiburcy et al. 2019). In contrast to the SMO approach, input cellularity in ESM can be optimally controlled. Day 22 myocytes (identified as optimal time point based on palpable expression of MRFs and ACTN2, **Figure 1E**) were dissociated and allowed to self- organize in the same collagen/Matrigel mixture as used for SMO generation. After formation of a compact tissue ring (after 4-6 days), ESM were transferred to metal holders for further maturation similar as described for the SMOs (**Figure 5A**). By 1-2 weeks maturation, spontaneous contractions were observed in ESM (**Video 2**) that increased in frequency until week 9 (**Video 3**). By 5 weeks of ESM maturation, compact muscle structure with parallel arrangement of myofibers could be observed. About 58±3% (n=3) of the cross-sectional area was populated with muscle cells embedded in a Laminin+ extracellular matrix (**Figure 5B**). Importantly, proteins of the DAG complex such as β-- dystroglycan, were properly localized to the cell membrane and demonstrated the compact arrangement of matured myofibers (**Figure 5C**). Ultrastructural analysis supported these findings showing advanced stages of myofibrillogenesis. Organized sarcomeres with distinct banding pattern including I- bands, A- bands, M-lines and Z disks were observed and mitochondria with dense matrix and developed cristae were found aligned with compact sarcomeres (**Figure 5D**). Average length of sarcomere in ESM was 1.8±0.02 µm (n=80). In addition, membranous structures of the triad, composed of a central T-tubule surrounded by two terminal cisternae from the sarcoplasmic reticulum were identified (Al-Qusairi and Laporte, 2011) (**Figure 5D**). Finally, we were interested if the ESM protocol is in principle suitable to also generate skeletal muscle from human primary skeletal myocytes (pSkM) isolated from 6 different patients with no known muscle disease. While force generating tissue was uniformly generated, we noticed high functional inter-patient variability in pSkM ESM compared to PSC ESM (**Supplementary Figure 2**).

**Figure 5.**
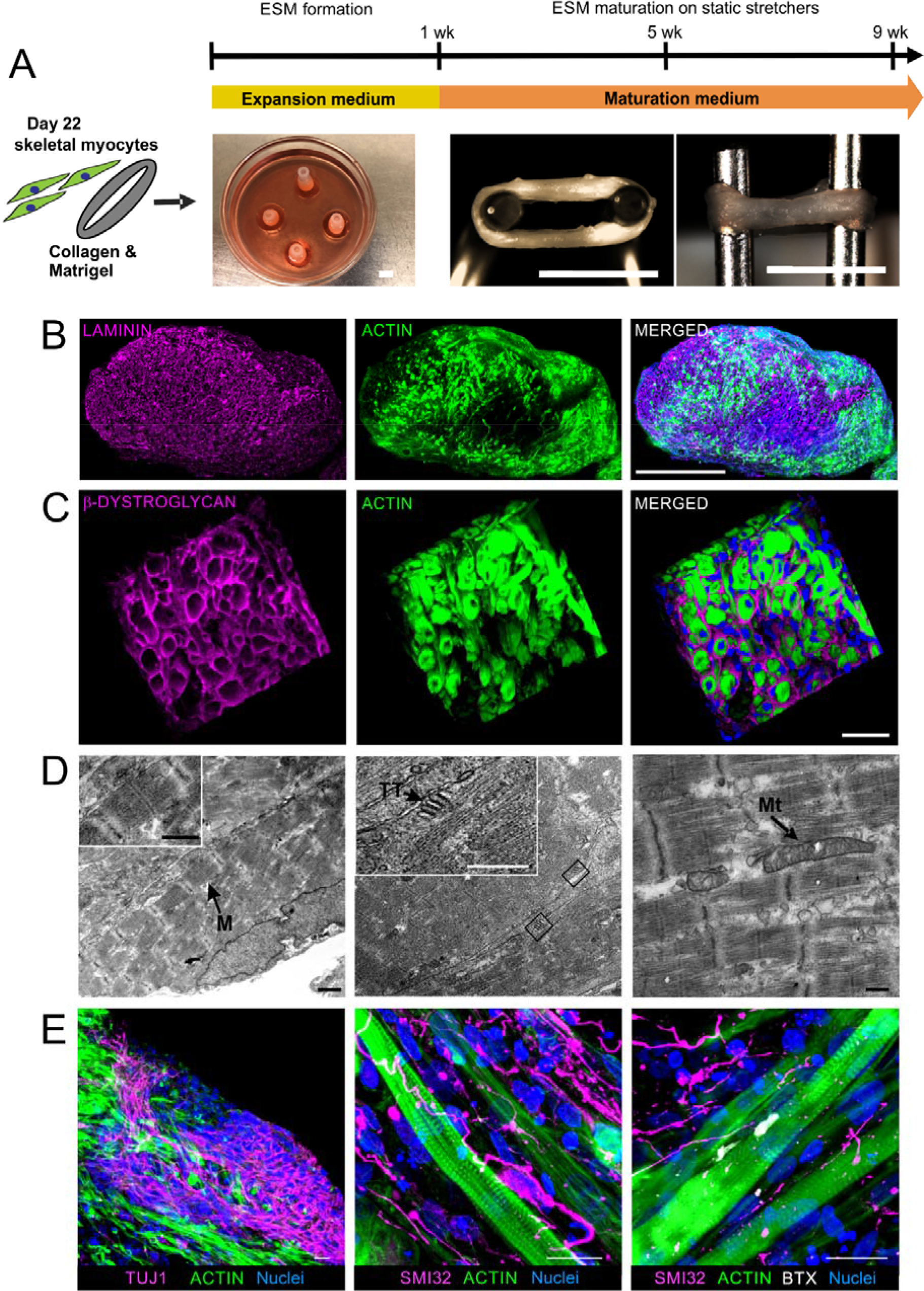
Advanced development of skeletal muscle structures in human engineered skeletal muscle. **(A)** Scheme of engineered skeletal muscle (ESM) generation from human PSC-derived skeletal myocytes with collagen type 1 and Matrigel™ in a ring-shaped hydrogel. ESM formation in expansion medium for 1 week in PDMS casting molds, functional maturation under isometric mechanical load (ESM on metal hooks of the static stretcher) for up to 9 wks. Scale bar: 5 mm. **(B)** Immunostaining of ACTIN+ muscle cells (green) and LAMININ+ extracellular matrix (magenta) in a representative cross section of 5 weeks old ESM. Scale bar: 500 µm. **(C)** Immunostaining of β-DYSTROGLYCAN (magenta) in the sarcolemma of ACTIN+ muscle fibers (green) in an ESM cross section. Scale bar: 40 µm. **(D)** Transmission electron microscopy (TEM) images of sarcomere ultrastructure, t-tubular triads and mitochondria along the muscle fibers in ESM. M: M-line, Mt: Mitochondria, TT: t-tubule. Scale bar: 1µm (left and middle panel) and 250 nm (right panel). **(E)** Immunostaining of ACTIN+ muscle cells (green) and TUJ1 or SMI32 positive neurons (magenta), Bungarotoxin+ (BTX, gray) motor end plates, and nuclei (blue) in 5 wks old ESM. Scale bars: 20 µm.

Another obvious difference between pSkM ESM and PSC ESM was the lack of spontaneous contractions in pSkM ESM. We therefore asked if this may be due to PSC ESM innervation by neurons. This assumption was based on the evidence of neuronal cells by single cell sequencing of the input day 22 skeletal myocyte cultures (**Figure 3C-F**) and neuronal differentiations in SMO (**Figure 4E**), but no evidence of the presence of neurons in pSkM preparations. To identify neurons in PSC skeletal myocyte preparations, we evaluated bulk RNA sequencing data of parallel day 60 monolayer cultures and day 60 ESM generated from the same day 22 cell source. Interestingly, we found a markedly higher abundance of neuronal transcripts (*ISL1, MNX1, LHX1*), suggestive for the presence of spinal cord motor neurons, in day 60 ESM compared to day 60 monolayer cultures (**Supplementary Figure 3A**). High abundance of *LHX1*, but not *LHX3*, does indeed suggest a motor neuron subpopulation with a limb muscle expression pattern (Sharma et al., 2000). The expression of mature neuronal and glial transcripts (*NSG2, GFAP*) was also significantly higher in ESM compared to 2D monolayer cultures at the same time point (day 60), suggesting generally more favorable conditions for neuronal co-development in ESM (**Supplementary Figure 3A)**. We further confirmed the presence of neuronal cell clusters, by immunostaining for neuron markers TUJ1 and SMI32. Interestingly, SMI32+ neurites seemed to contact α-bungarotoxin+ motor end plates on the muscle fibers, suggesting the self-organization of motor-end plate-like structures in ESM (**Figure 5E**). This was supported by expression of Muscle Associated Receptor Tyrosine Kinase (*MUSK*) which is critical for neuromuscular end plate organization (DeChiara et al., 1996) as well as acetylcholine choline esterase (*ACHE*) and nicotinic acetylcholine receptor subunit alpha (*CHRNA1*) (**Supplementary Figure 3B**). We confirmed the functionality of motor end plates by pharmacological activation of acetylcholine receptors. The mixed muscarinic-nicotinic acetylcholine receptor agonist carbachol caused a reversible depolarizing muscle block with almost complete ceasing of electrically stimulated contractions, indicating well-developed end plate function in ESM (**Supplementary Figure 3C,D)**.

### Maturation of myosin isoforms in ESM by T3 treatment

To enhance ESM maturation and based on the well documented role of thyroid hormones on muscle maturation as well as the finding of a high thyroid hormone receptor expression (blue “maturation cluster”; **Figure 2C**) we hypothesized that triiodo-L-thyronine (T3) addition may enhance the transition of myosin heavy chain isoform expression towards adult fast myosin isoforms (Larsson et al. 1994; Schiaffino et al. 1988, 2015; Simonides and van Hardeveld 2008) and increase tetanic force production. To test this hypothesis, we added T3 either during early maturation (1-5 weeks of ESM culture) or late maturation (5-9 weeks of ESM culture; **Figure 6A**).

**Figure 6:**
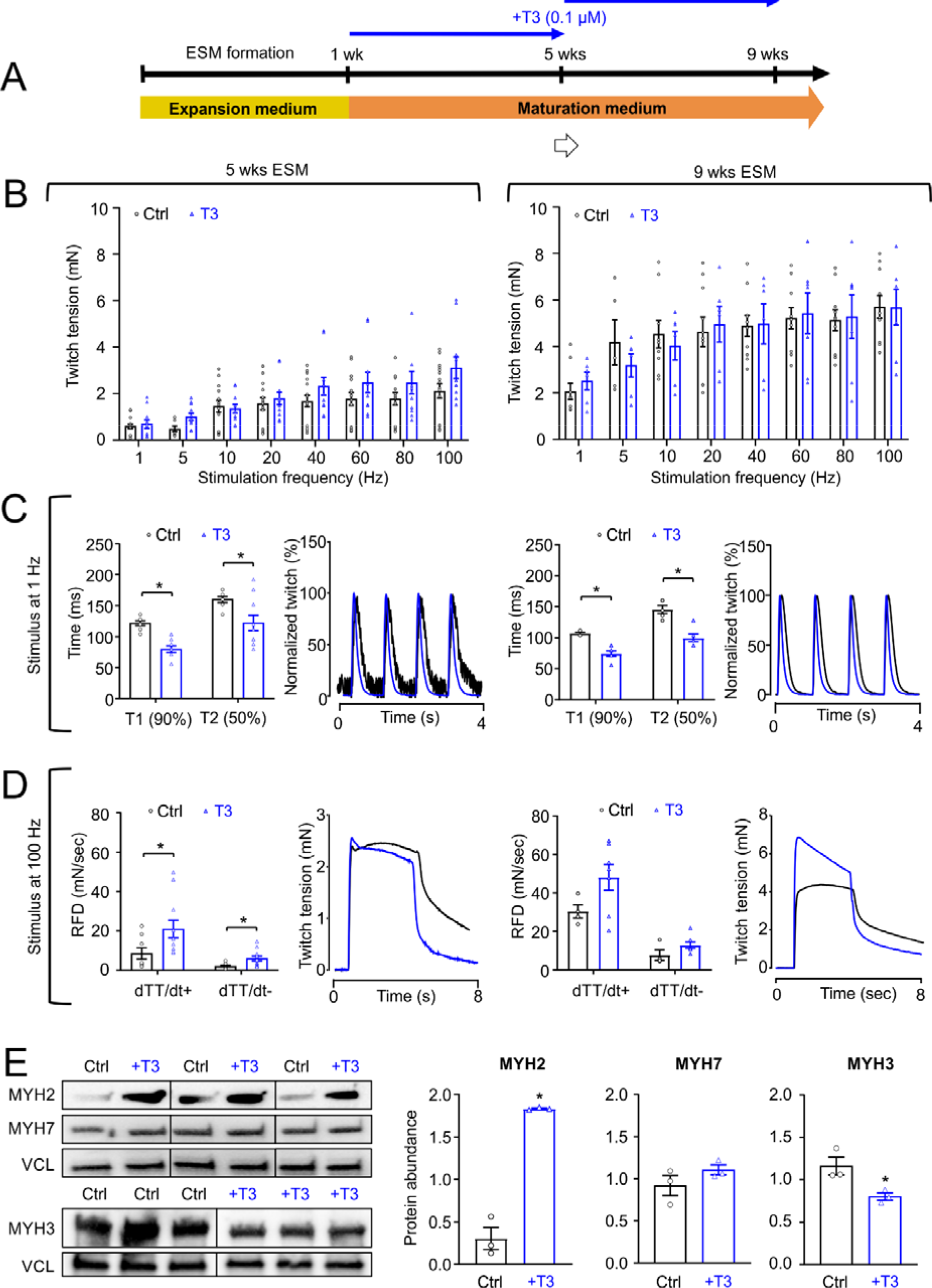
Advancing ESM function by thyroid hormone treatment. **(A)** Scheme of experimental design of ESM maturation for 5 or 9 wks with or without additional application of 0.1 µmol/L triiodo-L-thyronine (T3) for 4 wks. **(B)** Twitch tension in response to increasing stimulation frequencies of 5 wks and 9 wks old ESM cultured with (blue bars) or without T3 (black bars); n = 11-16/group at 5 wks; 7-10/group at 9 wks). **(C)** Quantification of contraction (T1) and relaxation (T2) time of single twitches of 5 wks old control (black bars) or +T3 (blue bars) ESM at 1 Hz **(first panel)**; Normalized representative traces of single twitches of 5 wks old control (black line) or +T3 (blue line) ESM at 1 Hz **(second panel)**; Quantification of contraction (T1) and relaxation (T2) time of single twitches of 9 wks old control (black bars) or +T3 (blue bars) ESM at 1 Hz **(third panel)**; Normalized representative traces of single twitches of 9 wks old control (black line) or +T3 (blue line) ESM at 1 Hz **(fourth panel)**; n = 5-11/group, *p<0.05 by Student’s t-test. (**D)** Rate of force development (RFD; rate of contraction: dTT/dt+ and rate of relaxation: dTT/dt-) of 5 wks old control (black bars) or +T3 (blue bars) ESM at 100 Hz tetanus **(first panel)**; Representative traces of twitch tension of 5 wks old control (black line) or +T3 (blue line) ESM at 100 Hz tetanus **(second panel)**; Rate of force development (RFD; rate of contraction: dTT/dt+ and rate of relaxation: dTT/dt-) of 9 wks old control (black bars) or +T3 (blue bars) ESM at 100 Hz tetanus **(third panel)**; Representative traces of twitch tension of 9 wks old control (black line) or +T3 (blue line) ESM at 100 Hz tetanus **(fourth panel)**; n = 4-11/group, *p<0.05 by Student’s t-test. **(E)** Immunoblot for fast myosin heavy chain (MYH2), slow myosin heavy chain (MYH7), embryonic myosin heavy chain (MYH3) and loading control vinculin (VCL). Protein abundance of MYH2 **(left panel)**, MYH7 **(middle panel)** and MYH3 **(right panel)** in 9 wks old ESM cultured with (blue bars) or without T3 (black bars); n = 3/group, *p<0.05 by Student’s t-test.

T3 treatment did not influence the maximal twitch tension (**Figure 6B**), but clearly shortened the duration of single twitches of both 5 and 9 week ESM. Accordingly, the speed of contraction (Time to 90% contraction - T1) of single twitches as well as relaxation (Time to 50% relaxation - T2) at 5 and 9 weeks was significantly increased (**Figure 6C**). In addition, the rate of force development (RFD) and the rate of force decline in tetanic contractions (100 Hz stimulation frequency) was enhanced in 5 weeks ESM and tended to be faster (p=0.18) also in 9 weeks ESM (**Figure 6D**).

The tetanus threshold (i.e. frequency where single twitches fuse to tetani) is greater in mammalian adult fast muscle fiber in comparison to slow muscle fibers (Buller and Lewis, 1965). The tetanus threshold of ESM with and without T3 treatment was analyzed by calculation of a fusion index (**Supplementary Figure 4**) derived from twitch recordings at increasing stimulation frequencies (**Figure 6B**). We found that the tetanus fusion index was different in ESM treated with T3 with a significant shift towards higher stimulation frequencies (50% fusion at 3.92±0.24 Hz vs 5.44±0.05 Hz in control ESM vs. ESM+T3, respectively, n=8; **Supplementary Figure 4**). Collectively, these functional data suggest that T3 enhances fast muscle properties of ESM.

We next asked if the T3 treatment affects the myosin heavy chain (MYH) isoform expression in ESM. Interestingly, T3 treatment clearly enhanced the abundance of mature fast MYH2 isoform with a reduction of the embryonic MYH3 isoform. The levels of the slow myosin isoform MYH7 were unchanged (**Figure 6E**). These molecular changes are well in line with the functional phenotype suggesting that T3 indeed supports maturation of fast skeletal muscle properties in ESM. The data also demonstrates that ESM respond to physiological stimuli comparable to skeletal muscle *in vivo*.

### ESM contain satellite cells with regenerative capacity

Even after prolonged *in vitro* culture we found muscle stem cell transcripts in the differentiated skeletal muscle cultures. In early (day 22) and late cultures (day 60), we found a similar expression of *PAX7,* but higher expression of satellite cell markers *MYF5* and *BARX2* (Cornelison and Wold, 1997; Meech et al., 2012) in ESM compared to monolayer culture suggesting a higher propensity to reconstitute and retain a satellite cell niche in ESM (**Figure 7A**). Immunostaining confirmed the presence of Pax7+ cells, 63±4% (n=267 cells counted) of which were located adjacent to a muscle fiber in ESM. The localization underneath the laminin+ basal lamina was indicative of a satellite cell position. Of note, 75±6% (n=164 cells counted) of these PAX7+ cells were Ki67-, implicating a quiescent state. In identically aged 2D monolayer cultures only 32±5% (n=105 cells counted) of PAX7+ cells were associated with muscle fibers (**Figure 7B**). These data suggest that muscle cells in ESM self-organize into myofibers with adjacent satellite cells and thus recapitulate important cellular components in an anatomically appropriate position for regenerative muscle.

**Figure 7.**
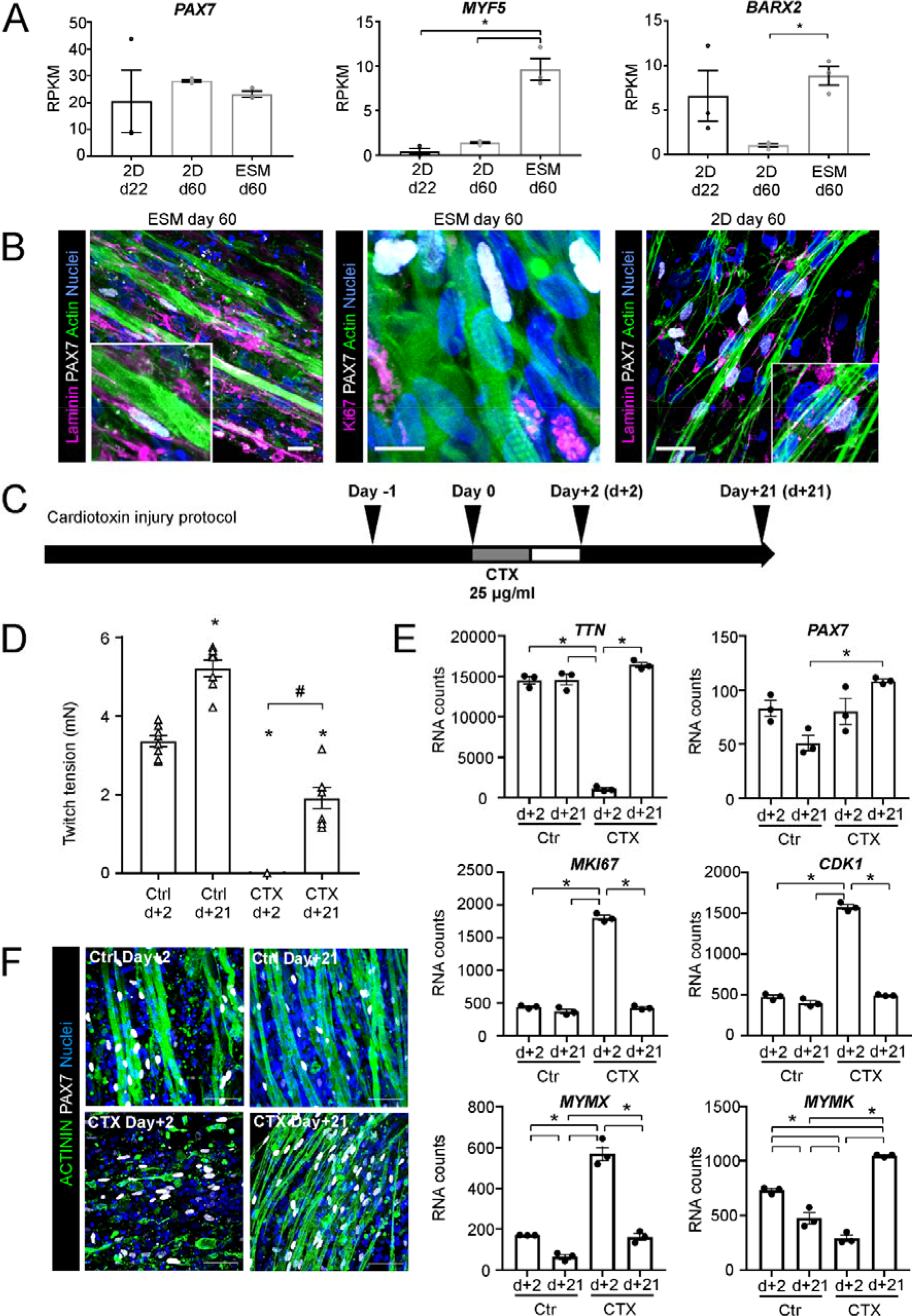
Regenerative capacity of human engineered skeletal muscle. **(A)** RNA transcript (Reads per Kilobase Million, RPKM) of indicated muscle stem cell markers in 2D monolayer cells at day 22 and day 60, plus day 60 ESM; n = 3-4/group, *p<0.05 by 1-way ANOVA and Tukey’s multiple comparison test. **(B)** Immunostaining of longitudinal sections of day 60 ESM for LAMININ (magenta), KI67 (magenta), ACTIN (green), PAX7 (gray), and nuclei (blue). Scale bars: 10 µm. Immunostaining of LAMININ (magenta), PAX7 (gray), ACTIN (green), and nuclei (blue) in 2D monolayer cultures at day 60. Scale bar: 50 µm. **(C)** Experimental design of cardiotoxin (CTX) injury model. ESM were incubated with 25 µg/ml CTX for 24 hrs. **(D)** Tetanic twitch tension at 100 Hz stimulation frequency of ESM at indicated time points after CTX (25 µg/ml) injury or control (Ctrl) condition; n=7-8/group, *p<0.05 vs. the respective Ctrl day +2, by 1-way ANOVA and Tukey’s multiple comparison test, #*p<0.05 CTX day +2 vs. CTX day +21. **(F)** RNA transcript for indicated genes at early (d+2) and late (day+21) time points after CTX (25 µg/ml) injury or control (Ctrl) conditions; n=3, *p<0.05 by 1-way ANOVA and Tukey’s multiple comparison test. **(F)** Immunostaining of sarcomeric a: -TININ (green), PAX7 (gray), and nuclei (blue) in ESM at indicated time points. Scale bars: 50 µm.

To test if these cells are capable of a muscle regeneration *in vitro,* we applied a cardiotoxin injury model in ESM [**Figure 7C**; (Tiburcy et al., 2019)]. 2 days after CTX injury ESM did not generate measurable force indicative of a complete loss of organized muscle fibers. After a regeneration period of 21 days, a partial, but robust recovery of contractile force (to 57±8% of initial force, n= 7) was observed (**Figure 7D**). RNA expression data were in line with the functional data showing an almost complete loss of mature muscle transcript (*TTN*) while *PAX7* transcript was largely preserved 2 days after CTX injury (**Figure 7E**). Upregulation of Ki67 (*MKI67*) and *CDK1* indicated cell cycle activation post injury. Consistent with results in murine muscle regeneration (Bi et al., 2017; Millay et al., 2014), we observed high expression of myomixer (*MYMX*) at day+2 coinciding with satellite cell activation. Myomaker (*MYMK*) expression increased later in the course of regeneration collectively showing that the regeneration of human muscle *in vitro* follows the regenerative pattern *in vivo*. Importantly, recovery of contractile force was paralleled by reexpression of *TTN* muscle transcript 21 days post injury. Immunostaining confirmed the almost complete loss of mature myofibers with sparing of PAX7+ satellite cells on day+2 after cardiotoxin injury. After 21 days of regeneration, substantial muscle was re-built (**Figure 7F**). To test if the cell cycle activation of satellite cells is required for ESM regeneration, we inhibited cell cycle activity by irradiation with 30 Gy. This completely abolished the regenerative response and formation of new muscle fibers (**Supplementary Figure 5**). Note, that irradiation of uninjured muscle did not impact contractile force. Those data demonstrate that ESM regeneration depends on activation of a PAX7+, dividing muscle stem cell.

## Discussion

We report a novel model for human skeletal muscle derivation in 2D and 3D organoid (SMO) cultures as well as for the engineering of skeletal muscle (ESM) with advanced structural and functional properties. Our data suggest that multicellularity (including neurons and supporting mesenchyme) as well as three-dimensionality are key for *in vitro* skeletal muscle development with *in vivo* properties. The re-engineering of a regeneration competent satellite cell niche appears particularly interesting as it may not only offer a solution for disease modelling and drug screening, but also for stable culture and amplification of satellite cells for regenerative applications (as demonstrated previously for the rat model; Tiburcy et al. 2019). The demonstration of further maturation of the developed limb muscle models under T3 supplementation further demonstrates a screening approach for maturation enhancing factors *in vitro*.

In contrast to MRF overexpression models (Albini et al., 2013; Darabi et al., 2012; Goudenege et al., 2012; Kim et al., 2017; Rao et al., 2018; Tedesco et al., 2012; Young et al., 2016), we directed differentiation in 2D-monolayer culture and 3D-organoids using defined and developmentally inspired growth factors and small molecules. The strength of this approach is the full control over the developmental origin of resulting muscle. We concentrated on generating hypaxial muscle as these muscle compartments (limbs, diaphragm and trunk) are predominantly affected by muscle diseases. Wnt activation (by CHIR99021), FGF signaling (FGF2), and BMP signaling inhibition (LDN193189) were instrumental in specifying paraxial mesoderm but not lateral plate mesoderm (increase in *MSGN1* but not *MESP1*). Following the induction of paraxial mesoderm, we demonstrated that maintaining FGF signaling in the presence Notch inhibition (DAPT) greatly increased the specification into somatic hypaxial progenitors. The predominant development into limb/trunk muscle was further supported by increase in transcript of migrating limb progenitors (*MEOX2, LBX1, MET*). Interestingly, generation of limb muscle progenitors was associated with *PITX2* and *MYOD1* expression but low levels of *MYF5* expression. This pattern fits to development of cells that first activate *MYOD1* expression to form limb muscles, whereas predominance of *MYF5* expression would indicate differentiation into “epaxial-like” muscles (Cossu et al., 1996; Kablar et al., 1997). In addition to the myogenic population, a *PDGFRA*+, *MEOX2*+, *EBF2*+ mesenchymal support cell population was identified that is reminiscent of limb mesenchyme (Reijntjes et al., 2007; Xi et al., 2020). It is conceivable that the mesenchymal population supports muscle formation also *in vitro*. Finally, we found evidence of co- developing neurons which expressed SMI32 contacting neuromuscular end plates. This was associated with expression of *MNX1, LHX1*, and *ISL1* but not *LHX3* supporting the development of motor neuron generation with a limb pattern (Sharma et al., 2000).

The co-development of spinal motor neurons has recently been described in PSC-derived skeletal muscle 3D organoids (Faustino Martins et al., 2020; Mazaleyrat et al., 2020). Interestingly, the data from our model share several aspects of the development of neuromuscular junctions by Faustino Martins et al. We observed efficient induction of neuromesodermal progenitor cells expressing *SOX2*, Brachyury (*T*) and *CDX2* as posterior axis “determinant” (Faustino Martins et al., 2020). Later cultures show significant expression of posterior HOXC genes 6,9, and 10 but not anterior axis genes (*FOXG1, OTX1, HOXB1*, not shown). Expression of “posterior axis” motor neuron genes *ISL1, MNX1* and *LHX1* increased between day 13 and 22 of differentiation at advanced stages of somitogenesis while expression of *OLIG2* as a marker of ventral spinal cord progenitors was not expressed. Collectively, we conclude that development of engineered skeletal muscle *in vitro* is associated with neuronal co-development of cells with high similarity to spinal cord motor neurons. The formation of neuromuscular junctions would likely increase maturation of ESM based on knowledge gained from previous studies in human 3D engineered muscle (Afshar Bakooshli et al., 2019).

While the SMO model allows for a simulation of embryonic muscle development and because of the simple one-step approach, our data demonstrates that more classical tissue engineering models, such as applied for the generation of ESM, are more likely to achieve higher levels of organotypic maturation. ESM demonstrated higher cellularity, clear anisotropic structure with membrane localized Dystrophin-associated protein complexes, advanced ultra-structural properties (e.g., Z- ,I- ,A- ,H- ,M-bands and t-tubulation) and ∼2-fold higher tetanic forces (2.3 vs 1.1 mN tetanic twitch force). The main difference is that the developmentally more advanced cellular input in ESM can be precisely controlled.

Despite the advanced organotypic properties, it is important to point out that the observed contractile parameters in ESM are not fully representative of adult skeletal muscle (Racca et al. 2013). For example, myocytes in ESM present with a smaller average muscle cell diameter (0.2-0.3 fold), a fetal myosin isoform expression pattern (high MYH3 to MYH2 ratio), ∼10% of the maximal contractile force reported for adult muscle, and high number of progenitor cells (PAX7+). Strategies to enhance physiological hypertrophic growth are needed to further enhance skeletal muscle properties. Increased MYH2 and reduced MYH3 expression under exposure to T3 represents first evidence of the propensity of ESM to undergo further maturation if exposed to a supportive environment. The fully serum-free process we established here will be advantageous for the testing of additional maturation factors.

Finally, the observation of regeneration in ESM after cardiotoxin-induced damage in dependence of PAX7+ satellite cell function was particularly notable because it demonstrates that regeneration-competent satellite cells can be developed using the reported protocol. Our conclusion that the observed PAX7+ in ESM are indeed satellite cells is based on the following observations: (i) PAX7-positive cells assume a satellite cells position underneath the basal lamina of muscle fibers; (ii) PAX7-positive cells are quiescent, but can be activated upon injury; (iii) irradiation blocks the regenerative response without evidence of functional impact on uninjured, for the most part post-mitotic ESM. The regeneration of engineered muscle by satellite cell activation is fascinating and has only recently been observed for human muscle *in vitro* (Fleming et al 2020). This study used primary muscle cells similar to earlier work in the rat (Juhas et al., 2014; Tiburcy et al., 2019) and a BaCl_2_ injury model, which may partially spare myotubes, leaving the possibility for PAX7+-cell-independent regeneration. To avoid this limitation, we have carefully titrated cardiotoxin to destroy most if not all developed myofibers, while sparing only and most of the PAX7+ cells. The following sequelae of PAX7+ cell activation, proliferation and fusion were completely inhibited by irradiation, which supports a true regenerative pattern.

We conclude that the skeletal muscle differentiation protocols in monolayer culture and in a novel organoid format (SMO) as well as the demonstration of skeletal muscle tissue engineering (ESM) as a means to enhance maturation provide novel platforms to study human hypaxial skeletal muscle development, disease and regeneration in a simple and robust *in vitro* model.

## Acknowledgments

M.T. is supported by the DZHK (German Center for Cardiovascular Research), and the German Research Foundation (DFG TI 956/1-1; SFB 1002 TP C04). W.H.Z. is supported by the DZHK (German Center for Cardiovascular Research), the German Federal Ministry for Science and Education (IndiHEART; 161L0250A), the German Research Foundation (DFG SFB 1002 C04/S01, IRTG 1816, MBExC) and the Fondation Leducq (20CVD04). Part of the work was supported by the Association Française contre les Myopathies (AFM, project no. 20987) to J.S. We acknowledge A.K. Hell and H.M. Lorenz for providing human skeletal muscle samples. Generation of the GMP line LiPSC-GR1.1 (also known as TC1133 or RUCDRi002-A) was supported by the NIH Common Fund Regenerative Medicine Program and reported in Stem Cell Reports (Baghbaderani et al. 2015). The NIH Common Fund and the National Center for Advancing Translational Sciences (NCATS) are joint stewards of the LiPSC-GR1.1 resource. The TC1133 line (Master Cell Bank Lot#: 50-001-21) was acquired by Repairon GmbH from the National Institute of Neurological Disorders and Stroke (NINDS) Human Cell and Data Repository (NHCDR) and processed to a GMP working cell bank (WCB). Post production cells from the WBC were kindly provided from Repairon GmbH to UMG for research use. Expert technical assistance by Iris Iben is gratefully acknowledged. J.Z. and J.S. are members of the European reference network for neuromuscular disorders (ERN EURO-NMD).

## Author contributions

Conceptualization, M.T., W-H.Z. and M.S.; Methodology, M.T., W-H.Z. and M.S.; Validation, M.T. and M.S.; Formal Analysis, M.T., M.S., M.R.I., M.S.S., D.K., H.A., O.S., M.Schm. and A.U.; Investigation, M.T., M.S., M.R.I., M.S.S., A.R., D.K., H.A., O.S., M.Schm. and A.U.; Resources, J.S. and J.Z.; Writing - Original Draft, M.T., M.S.,W-H.Z., M.R.I., M.S.S., O.S., J.Z., J.S. and A.U.; Writing - Review & Editing, M.T., W-H.Z. and M.S.; Visualization, M.T., M.S., M.R.I., M.S.S., A.R., D.K., L.K., H.A., O.S. and M.Schm.; Supervision, M.T., W-H.Z., A.F., W-A.L., G.S., J.S. and J.Z.; Funding Acquisition, M.T. and W-H.Z.

## EXPERIMENTAL PROCEDURES

### Human pluripotent stem cell culture

The following pluripotent stem cell (PSC) lines were used in the study: TC1133 [iPSC1; (Baghbaderani et al., 2015)], iPSC lines 2, 3, and 4 (Long et al., 2018), and HES2 (WiCell). The use of HES2 line was approved by the Robert-Koch-Institute (Nr. 3.04.02/0160). Informed consent and ethical approval by the University Medical Center Göttingen was obtained for use of human iPSC lines. All lines were routinely tested for pluripotency and confirmed to be free of mycoplasma (Lonza Mycoalert^TM^ kit). Human PSC lines were maintained on 1:120 Matrigel™ (BD) in phosphate-buffered saline (Thermo Fisher Scientific)–coated plates and cultured in StemMACS iPS-Brew XF (Miltenyi Biotec) at 37 °C and 5% CO_2_. Medium was changed every day and when the culture reached a confluency of 80-90%, it was rinsed once with PBS 1x (Thermo Fisher Scientific) and incubated in Versene solution (Thermo Fisher Scientific) for 3-5 min at room temperature. Versene was carefully aspirated and cells were gently washed off with StemMACS iPS-Brew XF (Miltenyi Biotec) with 5 μM Y27632 (Stemgent). Medium was changed to StemMACS iPS-Brew XF (Miltenyi Biotec) without Y27632 (Stemgent) after 24 hrs.

### Primary human myoblast culture

Muscle samples (Erector spinae muscle) were taken from patients during spine surgery after obtaining informed consent and with ethical approval by the University Medical Center Göttingen. Human muscle cell progenitors (satellite cells) were isolated according to the following protocol (Schmidt et al., 2008). In short, the muscle piece was minced and washed in phosphate buffered saline and trypsinized. The fragments were seeded to a 25-cm^2^ flask in skeletal muscle growth medium with supplement mix (PromoCell) and 1% Pen/Strep. After 21 days, myoblasts were labeled with anti-CD56/NCAM (mouse clone Eric-1; Thermo Fisher Scientific), followed by magnetic bead–labeled secondary antibodies and subsequently separated by magnets (Dynal/Invitrogen). For further expansion cells were seeded to T175 cell culture flasks in skeletal growth medium, which was replaced every other day.

### Directed differentiation of hPSCs into skeletal myocytes

Human pluripotent stem cells were plated at 1.3 x 10^4^ to 2.1 x 10^4^ cells/cm^2^ on 1:120 Matrigel™ (BD) in phosphate-buffered saline (Thermo Fisher Scientific) –coated plates and cultured in StemMACS iPS-Brew XF (Miltenyi Biotec) with 5 μ After 24 h, when the culture reached a confluency of 30 % (day 0), iPS-Brew XF was replaced with daily refreshed N2-CLF medium for 4 days. N2-CLF medium consisted of DMEM (Thermo Fisher Scientific) with 1% Pen/Strep, 1% N-2 Supplement, 1% MEM non-essential amino acid solution (all Thermo Fisher Scientific), 10 µM CHIR99021 (Stemgent), 0.5 µM LDN193189 (Stemgent) and 10 ng/ml FGF-2 (Peprotech). At this stage it is important to titrate cell density (colonies of ∼200 µm diameter) and CHIR concentration (7-10 µM) to prevent cell death and to perform medium changes slowly to avoid cell detachment. At day 4, the medium was exchanged with N2-FD medium every 24 hrs until day 6. N2-FD medium contained DMEM with 1% Pen/Strep), 1% N-2 Supplement, 1% MEM non-essential amino acid solution (all Thermo Fisher Scientific), 20 ng/ml FGF-2 (Peprotech) and 10 µM DAPT (Tocris). For day 6 and 7 the medium was replaced with N2-FDH medium which included DMEM with 1% Pen/Strep, 1% N-2 Supplement 1% MEM non-essential amino acid solution, 20 ng/ml FGF-2 (Peprotech), 10 µM DAPT (Tocris) and 10 ng/ml HGF (Peprotech). The medium was switched to N2-DHK medium on day 8, 9, 10 and 11. N2-DHK medium consisted of DMEM, with 1% Pen/Strep, 1% N-2 Supplement, 1% MEM non-essential amino acid solution (all Thermo Fisher Scientific), 10 µM DAPT (Tocris), 10 ng/ml HGF (Peprotech) and 10% knockout serum replacement (Thermo Fisher Scientific). From day 12 to 22, myogenic cells were cultured in expansion medium which was refreshed every second days. Expansion medium contained DMEM with 1% Pen/Strep, 1% N-2 Supplement, 1% MEM non-essential amino acid solution, 10% knockout serum replacement (all Thermo Fisher Scientific), and 10 ng/ml HGF (Peprotech). To further differentiate the cells to myotubes in monolayer culture, day 22 skeletal myocytes were enzymatically dissociated with TrypLE (Thermo Fisher Scientific) for 5 to 7 minutes at 37 °C and replated on 1:120 Matrigel™ (BD) in phosphate-buffered saline (Thermo Fisher Scientific)–coated plates at a density of 60–70,000 cells/cm^2^ in expansion medium with 5 μ M Y27632 (Stemgent). After 24 hr, the expansion medium was refreshed every other day for one week and then the medium was replaced with maturation medium. Maturation medium consisted of DMEM, with 1% Pen/Strep, 1% N-2 Supplement, and 2% B-27 Supplement (all Thermo Fisher Scientific). Maturation medium was changed every second day for 4 weeks.

### Cryopreservation of human PSC-derived skeletal myocytes

Human PSC-derived skeletal myocytes were cryopreserved on day 22 of culture. For enzymatic dissociation the cell culture was rinsed once with PBS 1x (Thermo Fisher Scientific). TrypLE (Thermo Fisher Scientific) was added to the cells and incubated for 5 to 7 minutes at 37 °C and 5% CO_2_. TrypLE digestion was stopped using expansion medium with 5 μM Y27632 (Stemgent). The cell suspension was triturated very gently with a 10-ml pipette to break the cell clumps and centrifuged at 100xg, 10 minutes, 21°C. Supernatant was removed and the pellet was resuspended very gently in freezing medium which contained μM Y27632 (Stemgent) and 10% DMSO (Sigma-Aldrich). 10x10^6^ human PSC-derived skeletal myocytes were frozen per cryovial in a MrFrosty™ freezing container (Nalgene) at -80°C overnight and then stored at -150°C.

### Thawing of human PSC-derived skeletal myocytes

The frozen cryovial was taken out from -150° deep freezer (SANYO) and quickly thawed in a water bath at 37° for approximately 2 min until a small ball of ice was still visible in the thawing medium. Using a 2 ml serological pipette, the contents of the cryovial were transferred to a pre-prepared 15 ml tube containing 9 ml of expansion medium with 5 μ Y27632 (Stemgent). The cell suspension was centrifuged at 100xg, 10 minutes, 21°C. The supernatant was removed and the pellet was resuspended very gently in expansion medium μM Y27632 (Stemgent) for downstream experiments.

### Preparation of casting molds and static stretchers

For the generation of the 3D muscle models, poly-dimethylsiloxane (PDMS; SYLGARDTM 184 Silicone Elastomer Kit, Dow Corning) circular molds with inner/outer diameter 4/6 mm and 2.5 mm height were fabricated and allowed to cure overnight at 55°C. Static stretch devices were made from a Teflon^®^ base and stainless steel holders. The detailed protocol for the preparation of the casting molds and static stretchers has been described previously (Soong et al., 2012; Tiburcy et al., 2014).

### Generation of human skeletal muscle organoids (SMOs)

To make skeletal muscle organoids (SMOs) from iPSCs, monolayer cultures were dissociated with Versene when reaching a confluency of 80-90%. A final 250 µl/SMO volume mixture of i) 0.23 mg acid soluble collagen type 1 (LLC Collagen Solutions), ii) 36 µl of concentrated 2x DMEM (Thermo Fisher Scientific) serum-free medium (0.27 g DMEM powder DMEM, powder, low glucose, pyruvate in 10 ml ddH2O), iii) 6.75 µl of NaOH 0.1 N (Carl Roth), iv) 10% v/v Matrigel™ (BD) and v) 0.8 x 10^6^ iPSC resuspended in 157.5 µl of StemMACS iPS-Brew XF (Miltenyi Biotec) medium with 5 μ(Peprotech) and 10% knockout serum replacement (ThermoFisher Scientific) was cast into circular PDMS molds (inner/outer diameter: 4/6 mm; height: 2.5 mm; volume: 250 μl). After 1 h of hydrogel polymerization at 37°C, StemMACS iPS-Brew XF (Miltenyi Biotec) medium with 5 μM Y27632 (Stemgent), 10 ng/ml FGF-2 (Peprotech) and 10% Knockout serum replacement (ThermoFisher Scientific) was added for 24 hrs. After tissue compaction skeletal muscle differentiation was induced following the protocol established in 2D. On day 22 of differentiation, SMOs were loaded on static stretchers at 120% of slack length and cultured in maturation medium for 4 additional weeks. Maturation medium was changed every second day and consisted of DMEM, low glucose, GlutaMAX™ Supplement, pyruvate (Thermo Fisher Scientific) with 1% Pen/Strep (Thermo Fisher Scientific), 1% N-2 Supplement (Thermo Fisher Scientific), 2% B-27 Supplement (Thermo Fisher Scientific) and 1 mM creatine monohydrate (Sigma-Aldrich).

### Generation of human engineered skeletal muscle

To generate human engineered skeletal muscle (ESM), either PSC-derived skeletal myocytes were dissociated, or frozen PSC-derived skeletal myocytes were thawed as described above. A final 250 µl/ESM volume mixture of i) 0.23 mg acid soluble collagen type 1 (Collagen Solutions), ii) 36 µl of concentrated 2x DMEM (Thermo Fisher Scientific) serum-free medium (0.27 g DMEM, powder, low glucose, pyruvate in 10 ml ddH2O), iii) 6.75 µl of NaOH 0.1 N (Carl Roth), iv) 10% v/v Matrigel™ (BD) and v) 1.25 x 10^6^ of day 22 hPSC-derived skeletal myocytes which is resuspended in 157.5 µl of expansion medium with 5 μ Y27632 (Stemgent), was cast into circular polydimethylsiloxane (PDMS) molds (inner/outer diameter: 4/6 mm; height: 2.5 mm; volume: 250 μl). After 1h of polymerization at 37°C, ESMs were cultured in expansion medium with 5 μY27632 (Stemgent) for 24 h and then expansion medium for another 6 days to consolidate into mechanically stable tissue. After transfer of ESMs onto static stretchers they were cultured in maturation medium under mechanical load up to 9 weeks. Maturation medium was changed every second day and consisted of DMEM with 1% Pen/Strep, 1% N-2 Supplement, 2% B-27 Supplement (all Thermo Fisher Scientific) and 0.1 µM T3 (Sigma-Aldrich). 1 mM creatine monohydrate (Sigma-Aldrich) was added to maturation medium from week 4 to 9.

ESM from primary skeletal myocytes was prepared in an identical way with the exception that cell resuspension and ESM culture was done in DMEM F12, 2 mmol/L L-glutamine, 15% FBS, 1% Pen Strep, 1:100 ITS-X supplement (all Thermo Scientific) for 48 hrs. ESM differentiation was then performed in DMEM with 1% Pen/Strep, 1% N-2 Supplement, 2% B- 27 Supplement (all Thermo Fisher Scientific).

### Isometric force measurements

Contractile function of ESM was measured under isometric conditions in a thermostatted organ bath (Föhr Medical Instruments) filled with gassed (5% CO_2_/95% O_2_) Tyrode’s solution (containing: 120 NaCl, 1 MgCl2, 0.2 CaCl2, 5.4 KCl, 22.6 NaHCO3, 4.2 NaH2PO4, 5.6 glucose, and 0.56 ascorbate; all in mmol/L) at 37°C. The calcium concentration was set to 1.8 mM. To normalize for the force-length relationship, ESMs were extended to Lmax (length of maximal twitch tension) under electrical stimulation with 1 Hz with 4 ms square pulses of 200 mA. At Lmax, twitch tension was assessed at varying frequencies (4-second long stimulation at 1,2,3,4,5, 10, 20, 40, 60, 80 and 100 Hz). At 1 Hz stimulation, depolarizing muscle block was induced by addition of the unspecific cholinergic receptor agonist carbachol (1 µmol/L). Contraction data was recorded with BMON software and analyzed using AMON software (Ingenieurbüro Jäckel).

### Cardiotoxin injury model

To induce muscle injury ESM was incubated with 25 µg/ml of of *Naja pallida* cardiotoxin (CTX; Latoxan) for 24 h in maturation medium (Tiburcy et al. 2019). Subsequently the injured tissue was rinsed and cultured in expansion medium for 1 week followed by maturation medium for another 2 weeks of regeneration. Medium was refreshed every second day. To irradiate the ESM prior to CTX injury the culture plate was placed in a STS Biobeam 8000 gamma irradiator and exposed to a single dose of 30 Gy irradiation over 10 minutes (Tiburcy et al., 2019).

### Immunostaining and confocal imaging

2D cell cultures were fixed in 4% formalin (Carl Roth) at 20-22°C for 15 min. ESM/BSM were fixed in 4% formalin at 4°C overnight. After 2 washes with PBS, ESM/BSM were plunged in 70% ethanol (Carl Roth) for 1 min and then embedded in 2% agarose (peqGOLD) in 1X Tris Acetate-EDTA (TAE) buffer. Using the Leica Vibrotome (LEICAVT1000S), sections were cut at 400 µm and kept in cold PBS. Prior to staining, 2D cell cultures as well as ESM sections were washed with PBS. For blocking and permeabilization, samples were incubated in blocking buffer (PBS with 5% fetal bovine serum, 1% bovine serum albumin (BSA), and 0.5% Triton-X). All the primary and secondary antibody staining was performed in the same blocking solution. The following antibodies were applied for primary staining at 20-22°C for 4h: Oct4 (1:500, Abcam), Pax3-concentrate (1:100, DSHB), Pax7-concentrate (1:100, DSHB), MyoD (1:100, Dako) and Myogenin-concentrate (1:10, DSHB), Sarcomeric α-actinin (1:500, Sigma-Aldrich), Laminin (1:50, Sigma-Aldrich), neurofilament H, SMI32 (1:20000, Biolegend), β-dystroglycan (1:50, LCL-b-DG, Leica Biosystem) and Ki67 (1:100, Abcam). After 3x PBS washes for 5 minutes, the appropriate Alexa Fluor-coupled secondary antibodies (1:1000, Thermo Fisher Scientific) were applied for 2h at 20-24°C. In parallel with secondary antibodies, Alexa Fluor 633-conjugated phalloidin (1:100, Thermo Fisher Scientific), Alexa Fluor 594-conjugated α Bungarotoxin and Hoechst 33342 (1:1000, Molecular Probes) were added to stain f-actin and nuclei, respectively. Following 3 washes with PBS, samples were mounted in Fluoromount-G (Southern Biotech). All the images were acquired by using a Zeiss LSM 710/NLO confocal microscope. To quantify the labeled cells, 3 randomly focal planes per sample from 3 different experiments were chosen for analysis with ImageJ cell counter tool.

### Transmission Electron Microscopy

Ultrastructural analysis was performed on ESM samples fixed in 4% formalin (Carl Roth), 15% saturated picric acid in 0.1 M PBS, pH 7.4, at 4°C overnight. ESMs were rinsed twice with PBS and treated with 0.5% OsO_4_ for 45 min following several washing steps in 100 mM phosphate buffer. Samples were counterstained with uranyl acetate, dehydrated via ethanol series, and embedded in Durcupan ACM epoxy resin (Sigma-Aldrich). Ultrathin sections were prepared from resin blocks using a Leica Ultracut S ultramicrotome (Mannheim, Germany) and adsorbed to glow-discharged formvar-carbon–coated copper single-slot grids. Electron micrographs were recorded using a Zeiss LEO 910 electron microscope; images were taken with a TRS Sharpeye CCD camera (Troendle, Moorenweis, Germany).

### Flow cytometry

Cells were fixed in 4% formalin (Carl Roth) at 20-22°C for 15 min. Following 2X washes with PBS, fixed samples were kept on ice for the staining process. Cells were incubated in blocking buffer (PBS with 5% fetal bovine serum, 1% bovine serum albumin (BSA), and 0.5% Triton-X) for 10 min. In parallel with staining for isotype controls, fixed cells were stained for Pax7 (1:500, DSHB), MyoD (1:500, Dako) and Myogenin (1:50, DSHB) and Sarcomeric α-actinin (1:4000, Sigma-Aldrich), for 45 min. Appropriate secondary antibodies (1:1000, Thermo Fisher Scientific) were applied for 30 min. Samples were stained with Hoechst-3342 for nuclear DNA counting and exclusion of cell doublets. Cells were run on a LSRII cytometer and at least 10,000 events per sample were analyzed using Diva software (BD Biosciences).

### Western blot analysis

For protein isolation snap frozen ESM was homogenized in 150 µl of ice-cold protein lysate buffer (2.38 g HEPES, 10.20 g NaCl, 100 ml Glycerol, 102 mg MgCl2, 93 mg EDTA, 19 mg EGTA, 5 ml NP-40 in a total volume of 500 ml ddH2O) containing phosphatase and protease inhibitor cocktail (Roche) then centrifuged for 30 min at 12000 rpm at 4°C. 30 µg of protein sample was loaded onto a 4 to 15% gradient sodium dodecyl sulfate (SDS)-polyacrylamid gel (Bio-Rad), followed by protein transfer to a polyvinylidene fluoride (PVDF) membrane. To visualize the total protein, the PVDF membrane was stained with Ponceau Red. Primary antibody (4 h at 20-22°C) and secondary antibody (1 h in at 20-22°C) staining was performed in blocking solution containing 5% milk in 1x Tris-buffered saline (TBS) and 0.1% Tween 20.

The following primary antibodies were applied: embryonic myosin heavy chain 3 (1:500, F1.652, DSHB), slow type myosin heavy chain 7 (1:500, A4.951, DSHB) and fast type myosin heavy chain 2 (1:100, A4.74, DSHB). Protein loading was controlled by Vinculin (VCL) antibody (1:5000, V3131, Sigma-Aldrich). Horseradish peroxidase conjugated goat anti-mouse IgG antibody (1:10000, P0260, Dako) was used for detection. The blots were developed with Femto LUCENT^TM^Luminol Reagent (Gbiosciences) and the protein bands were imaged using the BIO-RAD ChemDoc^TM^MP system. Protein quantification was performed using ImageJ.

### RNA expression analysis

RNA was purified using Trizol (Thermo Fisher Scientific) according to the manufacturer’s instructions and quantified using Nanodrop. To analyze skeletal muscle specific transcripts an nCounter Elements TagSet panel from was designed by NanoString Technologies. Fifty nanograms of RNA per sample were hybridized to target-specific capture and reporter probes at 67_°C overnight (20_h) according to the manufacturer’s instructions. Samples were then loaded into the NanoString cartridge and nCounter Gene Expression Assay started immediately. Raw reads were analyzed with nSolverTM Data Analysis Software. Background subtraction was performed using geometric means of negative controls. RNA counts were normalized to 4 housekeeping genes (*TBP, HPRT1, POL2RA, GAPDH)*.

### RNA sequencing

Prior to sequencing RNA quality was ensured with the Fragment Analyzer from Advanced Analytical by using the standard sensitivity RNA Analysis Kit (DNF-471). RNA-seq libraries were prepared using a modified strand-specific, massively-parallel cDNA sequencing (RNA-Seq) protocol from Illumina, the TruSeq Stranded Total RNA. Libraries were sequenced on a HiSeq 4000 platform (Illumina) generating 50 bp single-end reads (30-40 Mio reads/sample). Sequence images were transformed with Illumina software BaseCaller to BCL files, which was demultiplexed to fastq files with bcl2fastq v2.17.1.14. The quality check was done using FastQC (version 0.11.5). Sequence reads were aligned to the human genome reference assembly (UCSC version hg38) using Star (version 2.5.2a) (Dobin et al., 2013). For each gene, the number of mapped reads was counted for human genes in ENSEMBL annotation hg38 version 89 using featureCounts (version 1.4.5) (Liao et al., 2014). Raw counts were normalized and transformed to log2 counts per million (CPM) values. Reads Per Kilobase per Million mapped reads (RPKM) were calculated based on Ensembl transcript length using biomaRT (v2.24). All RNA sequencing data has been deposited in a public data base (GSE178270).

### Weighted co-expression analysis

Weighted gene co-expression network analysis was performed using (WGCNA) package (version 1.61) in R. Briefly, normalized counts were transformed into log (base 2) counts and were used to calculate pairwise bi-weighted mid-correlations between genes. Next, based on approximate scale-free topology a soft threshold power of 14 was chosen and was used to calculate pair-wise topological overlap between genes to construct a signed gene co-expression network. Modules of co-expressed genes was later identified based on following criteria: minimum module size of 100, method = “hybrid”, deepSplit =0, pamRespectsDendro=T, pamStage = T. Modules with correlation higher than 0.85 were merged together. Different modules were summarized as modular eigengenes (MEs), those were then used to compare expression of the given module across differentiation time points. The module specific genes were further filtered based on a module membership correlation coefficient cutoff of 0.60. Gene ontology of the modules were analyzed using clusterProfiler (v3.0.4) and after multiple adjustments only statistically significant gene ontology terms (FDR <0.05) were retrieved. For pathway analysis, Reactome (https://reactome.org/) database was used.

### Published dataset analysis

Raw data set from a previous study (Xi et al., 2017) was retrieved from NCBI GEO (accession: GSE90876) and processed as follows: sequencing reads were mapped to human genome hg38 using STAR aligner (v2.5.2b). After mapping, raw count files were generated using featureCounts of subread package (v1.5.1). For differential expression analysis, all samples were processed together and genes with less than 5 reads in 50% of the samples were filtered out prior to the analysis. Differential expression analysis was performed using DESeq2 package (version 1.28.1) in R. Genes with FDR < 0.05 were considered as differentially expressed. To test above chance overlap between previously identified module and differentially expressed genes, Fisheŕs exact test was performed.

### Single cell transcriptomics by single nuclei RNA sequencing

Single nuclei were isolated from flash frozen cells. The cell pellet was homogenized using a plastic pestle in a 1.5 ml Eppendorf tube containing 500 µl EZ prep lysis buffer (Sigma, NUC101-1KT) with 30 strokes. The homogenate was transferred into 2 ml microfuge tubes, lysis buffer was added up to 2 ml and incubated on ice for 7 minutes. After centrifuging for 5 minutes at 500xg supernatant was removed and the nuclear pellet was resuspended into 2 ml lysis buffer and incubated again on ice (7 minutes). After centrifuging for 5 minutes at 500xg, the supernatant was removed and the nuclei pellet was resuspended into 500 µl nuclei storage buffer (NSB: 1x PBS; Invitrogen, 0.5% RNase free BSA;Serva, 1:200 RNaseIN plus inhibitor; Promega, 1x EDTA-free protease inhibitor; Roche) and filtered through 40 μm filter (BD falcon) with additional 100 μL NSB to collect residual nuclei from the filter. Isolated nuclei were stained with a nuclear stain (7AAD) and FACS sorted (BD FACSaria III) to ensure a homogenous and viable nucleus preparation. Sorted nuclei were counted in a Countess FL II automated cell counter (ThermoFischer AMQAF1000, DAPI light cube: ThermoFischer: AMEP4650) with DAPI staining and nuclei concentration was adjusted to 1000 nuclei/μL. The nuclei were further diluted to capture and barcode 4000 nuclei according to Chromium single cell 3c reagent kit v3 (10X Genomics). Single nuclei barcoding, GEM formation, reverse transcription, cDNA synthesis and library preparation were performed according to 10X Genomics guidelines. cDNA libraries were pooled and sequenced 4 times in Illumina NextSeq 550 in order to achieve the target reads / nuclei. Each sequencing run was acquiring 150bp paired-end reads (Illumina NextSeq 550 High Output Kit v2.5). Demultiplexing, read mapping (to pre-mRNA reference genome) and gene counts per nuclei were computed with cellranger (v4.0) software. The nuclei barcoding and sequencing pipeline typically allows to obtain 50.000-100.000 reads/nucleus resulting in detection of 200-10.000 genes/nucleus (median: ∼2000 genes/nucleus) for further downstream analysis.

### Bioinformatic analysis of single-nucleus RNA-sequencing

Gene counts were obtained by aligning reads to the hg38 genome (NCBI:GCA 000001405.22) (GRCh38.p7) using CellRanger software (v.3.0.2) (10XGenomics). The CellRanger count pipeline was used to generate a gene-count matrix by mapping reads to the pre-mRNA as reference to account for unspliced nuclear transcripts. The SCANPY package was used for pre-filtering, normalization and clustering (Wolf et al., 2018). Initially, cells that reflected low-quality cells (based on read number and expression of house-keeping genes (Eisenberg and Levanon, 2013)) were excluded. Next, counts were scaled by the total library size multiplied by 10.000, and transformed to log space. Highly variable genes were identified based on dispersion and mean, the technical influence of the total number of counts was regressed out, and the values were rescaled. Principal component analysis (PCA) was performed on the variable genes, and UMAP was run on the top 50 principal components (PCs) (Becht et al., 2018). The top 50 PCs were used to build a k-nearest-neighbours cell–cell graph with k= 100 neighbors. Subsequently, spectral decomposition over the graph was performed with 50 components, and the Leiden graph-clustering algorithm was applied to identify cell clusters. We confirmed that the number of PCs captures almost all the variance of the data. For each cluster, we assigned a cell-type label using manual evaluation of gene expression for sets of known marker genes. A muscle gene panel was identified by calculating the differentially expressed genes between myogenic and non-muscle cluster with a low frequency cutoff of 1 and an adjusted p value of <0.05.

### Statistical analysis

All data were analyzed using GraphPad Prism 7 software (GraphPad Software Inc., San Diego) and presented as mean ±standard error of the mean (SEM). Statistical analyses were done using unpaired, two-tailed, Student’s t-test, 1-way or 2-way ANOVA where appropriate. Significantly different variances were corrected for. Results showing *p*<0.05 were considered significant and *n* indicates the number of samples.

## References

Afshar Bakooshli, M., Lippmann, E.S., Mulcahy, B., Iyer, N., Nguyen, C.T., Tung, K., Stewart, B.A., van den Dorpel, H., Fuehrmann, T., Shoichet, M., et al. (2019). A 3D culture model of innervated human skeletal muscle enables studies of the adult neuromuscular junction. eLife 8.

Al-Qusairi, L., and Laporte, J. (2011). T-tubule biogenesis and triad formation in skeletal muscle and implication in human diseases. Skelet Muscle 1, 26.

Albini, S., Coutinho, P., Malecova, B., Giordani, L., Savchenko, A., Forcales, S.V., and Puri, P.L. (2013). Epigenetic reprogramming of human embryonic stem cells into skeletal muscle cells and generation of contractile myospheres. Cell Rep 3, 661–670.

Baghbaderani, B.A., Tian, X., Neo, B.H., Burkall, A., Dimezzo, T., Sierra, G., Zeng, X., Warren, K., Kovarcik, D.P., Fellner, T., et al. (2015). cGMP-Manufactured Human Induced Pluripotent Stem Cells Are Available for Pre-clinical and Clinical Applications. Stem Cell Reports 5, 647–659.

Becht, E., McInnes, L., Healy, J., Dutertre, C.A., Kwok, I.W.H., Ng, L.G., Ginhoux, F., and Newell, E.W. (2018). Dimensionality reduction for visualizing single-cell data using UMAP. Nat Biotechnol.

Bi, P., Ramirez-Martinez, A., Li, H., Cannavino, J., McAnally, J.R., Shelton, J.M., Sanchez-Ortiz, E., Bassel-Duby, R., and Olson, E.N. (2017). Control of muscle formation by the fusogenic micropeptide myomixer. Science 356, 323–327.

Biressi, S., Molinaro, M., and Cossu, G. (2007). Cellular heterogeneity during vertebrate skeletal muscle development. Dev Biol 308, 281–293.

Bladt, F., Riethmacher, D., Isenmann, S., Aguzzi, A., and Birchmeier, C. (1995). Essential role for the c- met receptor in the migration of myogenic precursor cells into the limb bud. Nature 376, 768–771.

Borchin, B., Chen, J., and Barberi, T. (2013). Derivation and FACS-mediated purification of PAX3+/PAX7+ skeletal muscle precursors from human pluripotent stem cells. Stem Cell Reports 1, 620–631.

Buckingham, M. (2017). Gene regulatory networks and cell lineages that underlie the formation of skeletal muscle. Proc Natl Acad Sci U S A 114, 5830–5837.

Buckingham, M., and Mayeuf, A. (2012). Skeletal Muscle Development. Muscle Volume 2, 749–762.

Buller, A.J., and Lewis, D.M. (1965). The Rate of Tension Development in Isometric Tetanic Contractions of Mammalian Fast and Slow Skeletal Muscle. J Physiol 176, 337–354.

Caron, L., Kher, D., Lee, K.L., McKernan, R., Dumevska, B., Hidalgo, A., Li, J., Yang, H., Main, H., Ferri, G., et al. (2016). A Human Pluripotent Stem Cell Model of Facioscapulohumeral Muscular Dystrophy- Affected Skeletal Muscles. Stem Cells Transl Med 5, 1145–1161.

Chal, J., Al Tanoury, Z., Hestin, M., Gobert, B., Aivio, S., Hick, A., Cherrier, T., Nesmith, A.P., Parker, K.K., and Pourquie, O. (2016). Generation of human muscle fibers and satellite-like cells from human pluripotent stem cells in vitro. Nat Protoc 11, 1833–1850.

Chal, J., Oginuma, M., Al Tanoury, Z., Gobert, B., Sumara, O., Hick, A., Bousson, F., Zidouni, Y., Mursch, C., Moncuquet, P., et al. (2015). Differentiation of pluripotent stem cells to muscle fiber to model Duchenne muscular dystrophy. Nat Biotechnol 33, 962–969.

Choi, I.Y., Lim, H., Estrellas, K., Mula, J., Cohen, T.V., Zhang, Y., Donnelly, C.J., Richard, J.P., Kim, Y.J., Kim, H., et al. (2016). Concordant but Varied Phenotypes among Duchenne Muscular Dystrophy Patient-Specific Myoblasts Derived using a Human iPSC-Based Model. Cell Rep 15, 2301–2312.

Cornelison, D.D., and Wold, B.J. (1997). Single-cell analysis of regulatory gene expression in quiescent and activated mouse skeletal muscle satellite cells. Dev Biol 191, 270–283.

Cossu, G., Kelly, R., Tajbakhsh, S., Di Donna, S., Vivarelli, E., and Buckingham, M. (1996). Activation of different myogenic pathways: myf-5 is induced by the neural tube and MyoD by the dorsal ectoderm in mouse paraxial mesoderm. Development 122, 429–437.

Coumailleau, P., and Duprez, D. (2009). Sim1 and Sim2 expression during chick and mouse limb development. Int J Dev Biol 53, 149–157.

Darabi, R., Arpke, R.W., Irion, S., Dimos, J.T., Grskovic, M., Kyba, M., and Perlingeiro, R.C. (2012). Human ES- and iPS-derived myogenic progenitors restore DYSTROPHIN and improve contractility upon transplantation in dystrophic mice. Cell Stem Cell 10, 610–619.

Davis, R.L., Weintraub, H., and Lassar, A.B. (1987). Expression of a single transfected cDNA converts fibroblasts to myoblasts. Cell 51, 987–1000.

DeChiara, T.M., Bowen, D.C., Valenzuela, D.M., Simmons, M.V., Poueymirou, W.T., Thomas, S., Kinetz, E., Compton, D.L., Rojas, E., Park, J.S., et al. (1996). The receptor tyrosine kinase MuSK is required for neuromuscular junction formation in vivo. Cell 85, 501–512.

Dobin, A., Davis, C.A., Schlesinger, F., Drenkow, J., Zaleski, C., Jha, S., Batut, P., Chaisson, M., and Gingeras, T.R. (2013). STAR: ultrafast universal RNA-seq aligner. Bioinformatics 29, 15–21.

Eisenberg, E., and Levanon, E.Y. (2013). Human housekeeping genes, revisited. Trends Genet 29, 569–574.

Faustino Martins, J.M., Fischer, C., Urzi, A., Vidal, R., Kunz, S., Ruffault, P.L., Kabuss, L., Hube, I., Gazzerro, E., Birchmeier, C., et al. (2020). Self-Organizing 3D Human Trunk Neuromuscular Organoids. Cell Stem Cell 27, 498.

Goudenege, S., Lebel, C., Huot, N.B., Dufour, C., Fujii, I., Gekas, J., Rousseau, J., and Tremblay, J.P. (2012). Myoblasts derived from normal hESCs and dystrophic hiPSCs efficiently fuse with existing muscle fibers following transplantation. Mol Ther 20, 2153–2167.

Hirsinger, E., Malapert, P., Dubrulle, J., Delfini, M.C., Duprez, D., Henrique, D., Ish-Horowicz, D., and Pourquie, O. (2001). Notch signalling acts in postmitotic avian myogenic cells to control MyoD activation. Development 128, 107–116.

Juhas, M., Engelmayr, G.C., Jr., Fontanella, A.N., Palmer, G.M., and Bursac, N. (2014). Biomimetic engineered muscle with capacity for vascular integration and functional maturation in vivo. Proc Natl Acad Sci U S A 111, 5508–5513.

Kablar, B., Krastel, K., Ying, C., Asakura, A., Tapscott, S.J., and Rudnicki, M.A. (1997). MyoD and Myf-5 differentially regulate the development of limb versus trunk skeletal muscle. Development 124, 4729–4738.

Kim, J., Magli, A., Chan, S.S.K., Oliveira, V.K.P., Wu, J., Darabi, R., Kyba, M., and Perlingeiro, R.C.R. (2017). Expansion and Purification Are Critical for the Therapeutic Application of Pluripotent Stem Cell-Derived Myogenic Progenitors. Stem Cell Reports 9, 12–22.

Larsson, L., Li, X., Teresi, A., and Salviati, G. (1994). Effects of thyroid hormone on fast- and slow- twitch skeletal muscles in young and old rats. J Physiol 481 ( Pt 1), 149–161.

Lian, X., Hsiao, C., Wilson, G., Zhu, K., Hazeltine, L.B., Azarin, S.M., Raval, K.K., Zhang, J., Kamp, T.J., and Palecek, S.P. (2012). Robust cardiomyocyte differentiation from human pluripotent stem cells via temporal modulation of canonical Wnt signaling. Proc Natl Acad Sci U S A 109, E1848–1857.

Liao, Y., Smyth, G.K., and Shi, W. (2014). featureCounts: an efficient general purpose program for assigning sequence reads to genomic features. Bioinformatics 30, 923–930.

Long, C., Li, H., Tiburcy, M., Rodriguez-Caycedo, C., Kyrychenko, V., Zhou, H., Zhang, Y., Min, Y.L., Shelton, J.M., Mammen, P.P.A., et al. (2018). Correction of diverse muscular dystrophy mutations in human engineered heart muscle by single-site genome editing. Sci Adv 4, eaap9004.

Maffioletti, S.M., Sarcar, S., Henderson, A.B.H., Mannhardt, I., Pinton, L., Moyle, L.A., Steele-Stallard, H., Cappellari, O., Wells, K.E., Ferrari, G., et al. (2018). Three-Dimensional Human iPSC-Derived Artificial Skeletal Muscles Model Muscular Dystrophies and Enable Multilineage Tissue Engineering. Cell Rep 23, 899–908.

Mamchaoui, K., Trollet, C., Bigot, A., Negroni, E., Chaouch, S., Wolff, A., Kandalla, P.K., Marie, S., Di Santo, J., St Guily, J.L., et al. (2011). Immortalized pathological human myoblasts: towards a universal tool for the study of neuromuscular disorders. Skelet Muscle 1, 34.

Mayeuf-Louchart, A., Lagha, M., Danckaert, A., Rocancourt, D., Relaix, F., Vincent, S.D., and Buckingham, M. (2014). Notch regulation of myogenic versus endothelial fates of cells that migrate from the somite to the limb. Proc Natl Acad Sci U S A 111, 8844–8849.

Mazaleyrat, K., Badja, C., Broucqsault, N., Chevalier, R., Laberthonniere, C., Dion, C., Baldasseroni, L., El-Yazidi, C., Thomas, M., Bachelier, R., et al. (2020). Multilineage Differentiation for Formation of Innervated Skeletal Muscle Fibers from Healthy and Diseased Human Pluripotent Stem Cells. Cells 9.

Meech, R., Gonzalez, K.N., Barro, M., Gromova, A., Zhuang, L., Hulin, J.A., and Makarenkova, H.P. (2012). Barx2 is expressed in satellite cells and is required for normal muscle growth and regeneration. Stem Cells 30, 253–265.

Mendjan, S., Mascetti, V.L., Ortmann, D., Ortiz, M., Karjosukarso, D.W., Ng, Y., Moreau, T., and Pedersen, R.A. (2014). NANOG and CDX2 pattern distinct subtypes of human mesoderm during exit from pluripotency. Cell Stem Cell 15, 310–325.

Millay, D.P., Sutherland, L.B., Bassel-Duby, R., and Olson, E.N. (2014). Myomaker is essential for muscle regeneration. Genes Dev 28, 1641–1646.

Miura, S., Davis, S., Klingensmith, J., and Mishina, Y. (2006). BMP signaling in the epiblast is required for proper recruitment of the prospective paraxial mesoderm and development of the somites. Development 133, 3767–3775.

Rajakumari, S., Wu, J., Ishibashi, J., Lim, H.W., Giang, A.H., Won, K.J., Reed, R.R., and Seale, P. (2013). EBF2 determines and maintains brown adipocyte identity. Cell Metab 17, 562–574.

Rao, L., Qian, Y., Khodabukus, A., Ribar, T., and Bursac, N. (2018). Engineering human pluripotent stem cells into a functional skeletal muscle tissue. Nat Commun 9, 126.

Reijntjes, S., Stricker, S., and Mankoo, B.S. (2007). A comparative analysis of Meox1 and Meox2 in the developing somites and limbs of the chick embryo. Int J Dev Biol 51, 753–759.

Rios, A.C., Serralbo, O., Salgado, D., and Marcelle, C. (2011). Neural crest regulates myogenesis through the transient activation of NOTCH. Nature 473, 532–535.

Schiaffino, S., Ausoni, S., Gorza, L., Saggin, L., Gundersen, K., and Lomo, T. (1988). Myosin heavy chain isoforms and velocity of shortening of type 2 skeletal muscle fibres. Acta Physiol Scand 134, 575–576.

Schmidt, J., Barthel, K., Wrede, A., Salajegheh, M., Bahr, M., and Dalakas, M.C. (2008). Interrelation of inflammation and APP in sIBM: IL-1 beta induces accumulation of beta-amyloid in skeletal muscle. Brain 131, 1228–1240.

Selvaraj, S., Mondragon-Gonzalez, R., Xu, B., Magli, A., Kim, H., Laine, J., Kiley, J., McKee, H., Rinaldi, F., Aho, J., et al. (2019). Screening identifies small molecules that enhance the maturation of human pluripotent stem cell-derived myotubes. eLife 8.

Sharma, K., Leonard, A.E., Lettieri, K., and Pfaff, S.L. (2000). Genetic and epigenetic mechanisms contribute to motor neuron pathfinding. Nature 406, 515–519.

Shelton, M., Kocharyan, A., Liu, J., Skerjanc, I.S., and Stanford, W.L. (2016). Robust generation and expansion of skeletal muscle progenitors and myocytes from human pluripotent stem cells. Methods 101, 73–84.

Shelton, M., Metz, J., Liu, J., Carpenedo, R.L., Demers, S.P., Stanford, W.L., and Skerjanc, I.S. (2014). Derivation and expansion of PAX7-positive muscle progenitors from human and mouse embryonic stem cells. Stem Cell Reports 3, 516–529.

Simonides, W.S., and van Hardeveld, C. (2008). Thyroid hormone as a determinant of metabolic and contractile phenotype of skeletal muscle. Thyroid 18, 205–216.

Soong, P.L., Tiburcy, M., and Zimmermann, W.H. (2012). Cardiac differentiation of human embryonic stem cells and their assembly into engineered heart muscle. Curr Protoc Cell Biol Chapter 23, Unit23 28.

Striedinger, K., Barruet, E., and Pomerantz, J.H. (2021). Purification and preservation of satellite cells from human skeletal muscle. STAR Protoc 2, 100302.

Tedesco, F.S., Gerli, M.F., Perani, L., Benedetti, S., Ungaro, F., Cassano, M., Antonini, S., Tagliafico, E., Artusi, V., Longa, E., et al. (2012). Transplantation of genetically corrected human iPSC-derived progenitors in mice with limb-girdle muscular dystrophy. Sci Transl Med 4, 140ra189.

Tiburcy, M., Hudson, J.E., Balfanz, P., Schlick, S., Meyer, T., Chang Liao, M.L., Levent, E., Raad, F., Zeidler, S., Wingender, E., et al. (2017). Defined Engineered Human Myocardium With Advanced Maturation for Applications in Heart Failure Modeling and Repair. Circulation 135, 1832–1847.

Tiburcy, M., Markov, A., Kraemer, L.K., Christalla, P., Rave-Fraenk, M., Fischer, H.J., Reichardt, H.M., and Zimmermann, W.H. (2019). Regeneration competent satellite cell niches in rat engineered skeletal muscle. FASEB Bioadv 1, 731–746.

Tiburcy, M., Meyer, T., Soong, P.L., and Zimmermann, W.H. (2014). Collagen-based engineered heart muscle. Methods Mol Biol 1181, 167–176.

Uezumi, A., Fukada, S., Yamamoto, N., Takeda, S., and Tsuchida, K. (2010). Mesenchymal progenitors distinct from satellite cells contribute to ectopic fat cell formation in skeletal muscle. Nat Cell Biol 12, 143–152.

Wolf, F.A., Angerer, P., and Theis, F.J. (2018). SCANPY: large-scale single-cell gene expression data analysis. Genome Biol 19, 15.

Xi, H., Fujiwara, W., Gonzalez, K., Jan, M., Liebscher, S., Van Handel, B., Schenke-Layland, K., and Pyle, A.D. (2017). In Vivo Human Somitogenesis Guides Somite Development from hPSCs. Cell Rep 18, 1573–1585.

Xi, H., Langerman, J., Sabri, S., Chien, P., Young, C.S., Younesi, S., Hicks, M., Gonzalez, K., Fujiwara, W., Marzi, J., et al. (2020). A Human Skeletal Muscle Atlas Identifies the Trajectories of Stem and Progenitor Cells across Development and from Human Pluripotent Stem Cells. Cell Stem Cell.

Xu, B., Zhang, M., Perlingeiro, R.C.R., and Shen, W. (2019). Skeletal Muscle Constructs Engineered from Human Embryonic Stem Cell Derived Myogenic Progenitors Exhibit Enhanced Contractile Forces When Differentiated in a Medium Containing EGM-2 Supplements. Adv Biosyst 3, e1900005.

Young, C.S., Hicks, M.R., Ermolova, N.V., Nakano, H., Jan, M., Younesi, S., Karumbayaram, S., Kumagai-Cresse, C., Wang, D., Zack, J.A., et al. (2016). A Single CRISPR-Cas9 Deletion Strategy that Targets the Majority of DMD Patients Restores Dystrophin Function in hiPSC-Derived Muscle Cells. Cell Stem Cell 18, 533–540.

