## Supplementary material for "Human engineered skeletal muscle of hypaxial origin from pluripotent stem cells with advanced function and regenerative capacity"

**Supplementary table 1 related to Figure 2: Biological process annotation of coexpression clusters.**

| Cluster identifier | Biological process annotated | Enriched GO terms |
|---|---|---|
| Black | Migrating limb progenitors | Muscle organ development |
| Blue | Muscle maturation | Regulation of neuron development; Muscle contraction |
| Brown | Presomitic progenitor development | Regulation of cell cycle phase transition |
| Cyan | None | No enrichment |
| Darkgreen | Cellular respiration | Oxidative phosphorylation |
| Darkred | Myoblast development | Hexose metabolic process |
| Green | Cilium organization | Cilium organization |
| Greenyellow | Histone modification | Histone modification |
| Grey | Primitive streak development | Mitochondrial gene expression |
| Lightcyan | DNA organization | DNA conformation change |
| Lightgreen | Unfolded protein response | Autophagy; Response to unfolded protein |
| Lightyellow | None | No enrichment |
| Magenta | Translation | Ribonucleoprotein complex biogenesis |
| Midnightblue | Translation | ncRNA metabolic process |
| Pink | Myotube development | Response to endoplasmic reticulum stress |
| Purple | None | No enrichment |
| Red | Presomitic progenitor development | DNA replication |
| Royalblue | Lipid storage | Lipid storage |
| Salmon | Dermomyotome development | Muscle tissue development |
| Tan | None | No enrichment |
| Turquoise | Somitic progenitor development | Cilium organization |
| Yellow | Pluripotency | Telomere maintenance via telomerase |

### Supplementary Figure 1

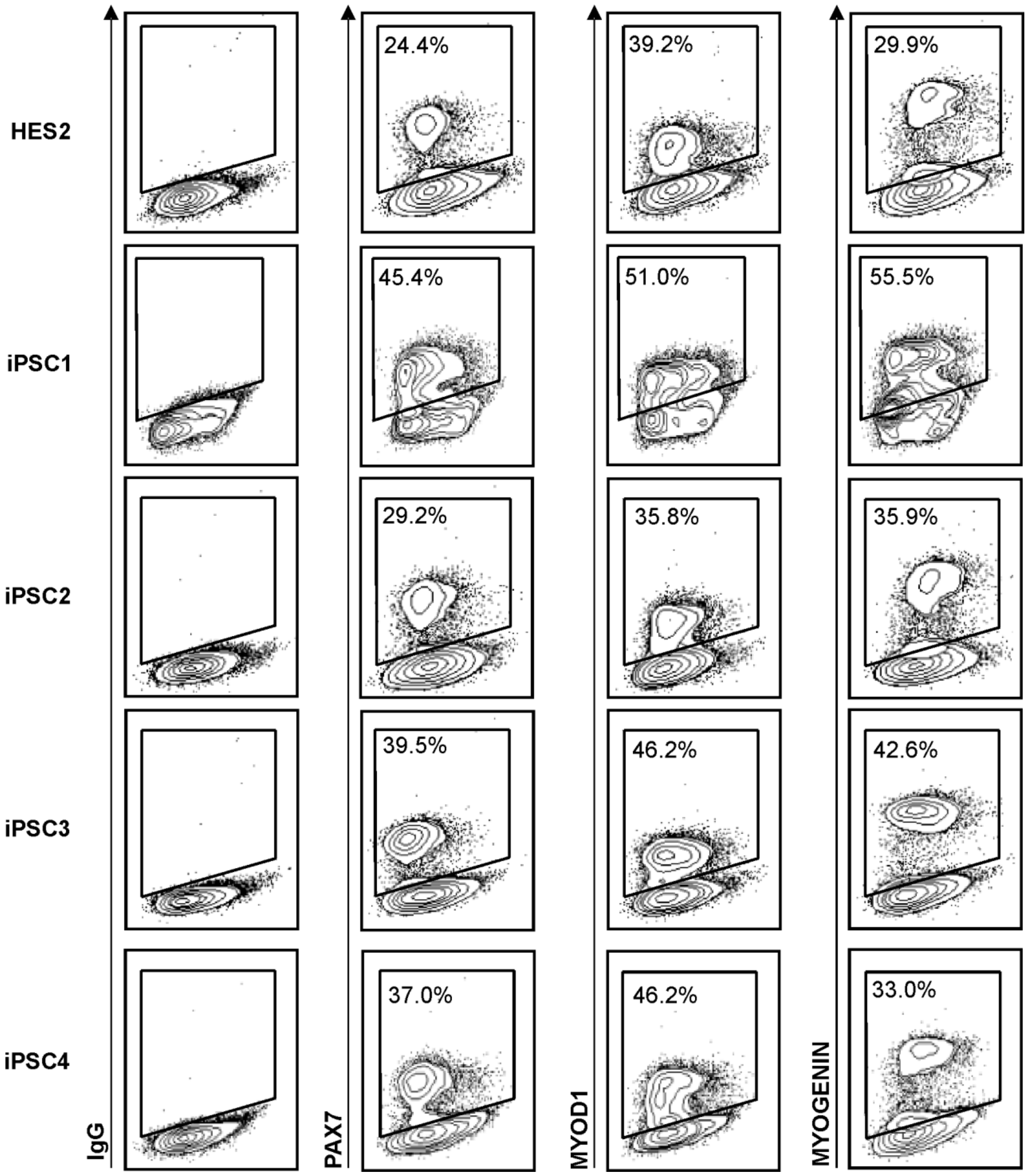


**Supplementary Figure 1 related to Figure 3: Efficiency of skeletal myocyte differentiation from human PSC**

Flow cytometry of myogenic regulatory factors PAX7, MYOD1, MYOGENIN in comparison to isotype control (IgG) in day 22 old skeletal myocyte cultures from different PSC lines.

**Supplementary Table 2 related to Figure 3**

**List of muscle enriched genes (muscle_gene panel)**

*ABCC9, ABCG1, ABLIM3, ACADL, ACOT11, ACTC1, ACTN2, ACVR2A, ADAM2, ADAMTS20, ADAMTS5, ADGRA1, AFAP1L1, AKR1C1, AKR1C2, AKR1C3, AL138752.2, AL591806.4, ALPK2, ALPK3, AP006333.1, ARHGAP18, ARHGAP9, ARMC3, ARPP21, ASB5, ATP2A1, ATP2B1, B4GALNT3, BBOX1, BCL11B, BICDL1, BLCAP, BMPR1B, BVES, C10ORF90, C1QTNF7, CACNA1S, CACNA2D1, CACNA2D4, CAP2, CASQ2, CASS4, CASZ1, CCDC141, CCNG2, CD82, CD96, CDH15, CDH7, CDK15, CELF2, CFL2, CHD7, CHODL, CHRNA1, CHRNB1, CHRND, CHRNG, CLCN5, CLSTN2, COBL, COL19A1, COL25A1, COQ8A, CPM, CYB5R1, DACH1, DES, DGKB, DLG2, DNAH17, DOCK6, DOCK9, DOK7, DSCAML1, DUSP10, DUSP27, DYSF, ECHDC2, EFHD2, EHBP1L1, EMC10, ENO3, ENPP1, ERBB3, EYA1, EYA2, EYA4, EZR, FASTKD1, FAT1, FGD4, FGF10, FGF13, FGF9, FGFR4, FNDC5, FOXO1, FOXP2, FREM2, FRMD3, FRMPD1, FST, FSTL4, GABRB3, GADL1, GATM, GCNT1, GEN1, GLI1, GLRB, GPA33, GPRIN3, GRAMD1B, GREM2, GSG1L, HES, HEYL, HIPK4, HS6ST2, IGDCC4, INPP4, ITGA4, ITGA7, ITGB6, ITIH5, JAM3, JPH1, JPH2, JSRP1, KCND3, KCNN2, KIF24, KLHL13, KLHL14, KLHL31, KLHL41, KREMEN1, KREMEN2, LDB3, LDLRAD3, LFNG, LHFPL6, LINC00514, LMOD3, LRIG1, LRP5, LRRC3B, LRRFIP1, LZTS1, MACROD1, MAMSTR, MAN1C1, MAP4K1, MARCHF3, MATR3_2, MB21D2, MEF2C, MEGF10, MET, MICAL1, MICAL2, MLIP, MMP23B, MRLN, MSR1, MTHFD1L, MYH3, MYL1, MYL4, MYLK4, MYO18B, MYOG, MYOM3, MYOZ2, MYPN, NCOA1, NEB, NECTIN1, NES, NEXN, NKAIN4, NNAT, NPNT, NRK, NTF3, NTN5, NXPH2, OLFML2A, OLFML2B, ORC4, OVCH1, P3H2, PALM2AKAP2, PALMD, PARM1, PAX7, PC, PDE1C, PDGFC, PDLIM3, PITPNM3, PITX2, PITX3, PKP4, PLAC1, PLS3, PLXNA2, POLA1, PPFIA4, PRELID3A, PRKCB, PRUNE2, PSEN2, PTGFR, PUS7, RALYL, RAPSN, RASSF3, RASSF4, RBM20, RBM24, RCL1, RELL1, RGS7, RIF1, RYR1, RYR2, SCN7A, SEMA3D, SEMA6B, SEPTIN4, SETD7, SGCA, SGCD, SHD, SHISA9, SIM1, SKP2, SLC16A10, SLC24A2, SLC24A3, SLC38A5, SLC7A2, SLC8A3, SLF2, SMC6, SMOC1, SMYD1, SNTG2, SOX6, SPAG6, SPATS2L, SRL, SRPK3, ST6GALNAC5, ST7, STARD13, STC1, STC2, STK26, SYN2, SYNE3, SYTL3, TANC1, TEAD4, TMEM131L, TMEM232, TMTC1, TNC, TNNI1, TNNT1, TNNT2, TNNT3, TNPO1, TPM2, TRDN, TRIM55, TRIM72, TRPA1, TSHZ3, TSPAN12, TSPAN33, UNC45B, USP6, VGLL2, VGLL3, VWCE, WDR43, ZEB2, ZNF536*

### Supplementary Figure 2

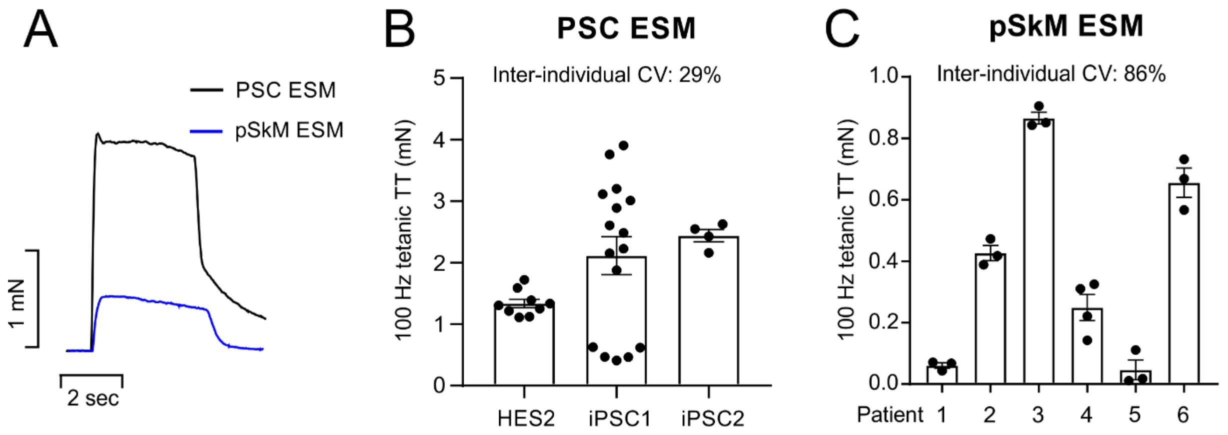


**Supplementary Figure 2 related to Figure 5. Functional variability of ESM from PSC and biopsy-derived skeletal myocytes. (A)** Representative original twitch tension traces at 100 Hz tetanic contraction of ESM prepared from PSC-derived (PSC ESM) or biopsy-derived primary SkM (pSkM ESM). **(B)** Tetanic twitch tension (TT) at 100 Hz of ESM from one HES (HES2) and two wildtype iPSC lines. The inter-individual coefficient of variation (CV) is indicated. **(C)** Tetanic twitch tension (TT) at 100 Hz of ESM from 6 different patient biopsies. After CD56 purification skeletal myoblasts were expanded for an average 70±16 days (n=5). The inter-individual coefficient of variation (CV) is indicated.

### Supplementary Figure 3

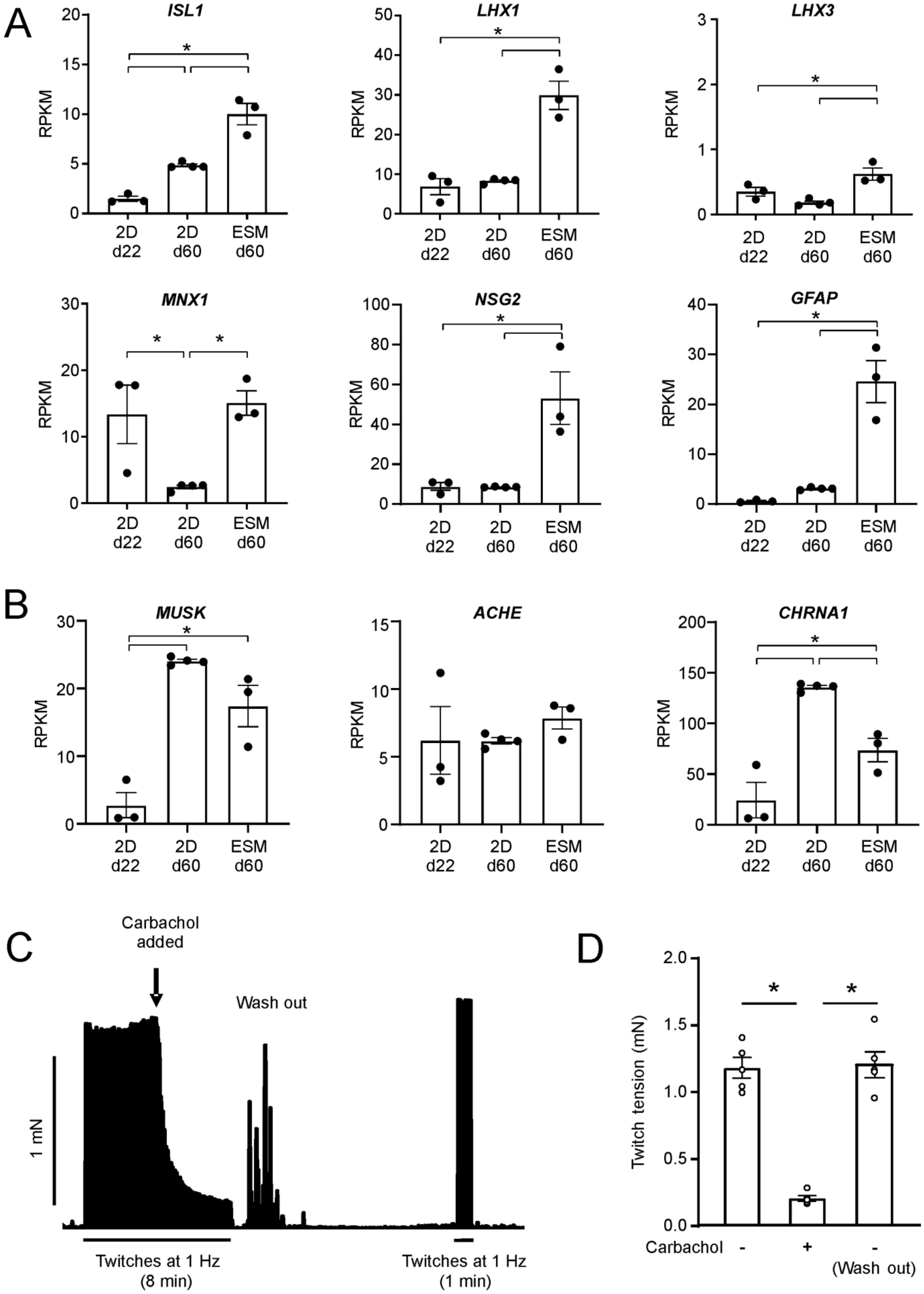

**Supplementary Figure 3 related to Figure 5. Neuronal co-development in engineered skeletal muscle. (A)** RNA transcript (Reads per Kilobase Million, RPKM) of neuronal markers in source 2D monolayer cells at day 22 and parallel cultures of 2D monolayer at day 60 and ESM day 60; n = 3-4/group, *$p<0.05$ by 1-way ANOVA and Tukey's multiple comparison test. **(B)** RNA transcript (Reads per Kilobase Million, RPKM) of motor end plate markers in source 2D monolayer cells at day 22 and parallel cultures of 2D monolayer at day 60 and ESM day 60; n = 3-4/group, *$p<0.05$ by 1-way ANOVA and Tukey's multiple comparison test. Immunostaining of ACTIN+ muscle cells (green) and TUJ1 or SMI32 positive neurons (magenta), Bungarotoxin+ (BTX, gray) motor end plates, and nuclei (blue) in 5 wks old ESM. Scale bars: 20 µm **(C)** Representative recording of ESM twitch tension (bar indicates 1 mN) at 1 Hz electrical stimulation. The unspecific cholinergic receptor agonist carbachol (1 µM) is added where indicated and later washed out from the organ bath. **(D)** Quantification of twitch tension at 1 Hz stimulation before and after treatment of ESM with 1 µM carbachol and after washout; n = 5; *$p<0.05$ by 1-way ANOVA and Tukey's multiple comparison test.

### Supplementary Figure 4

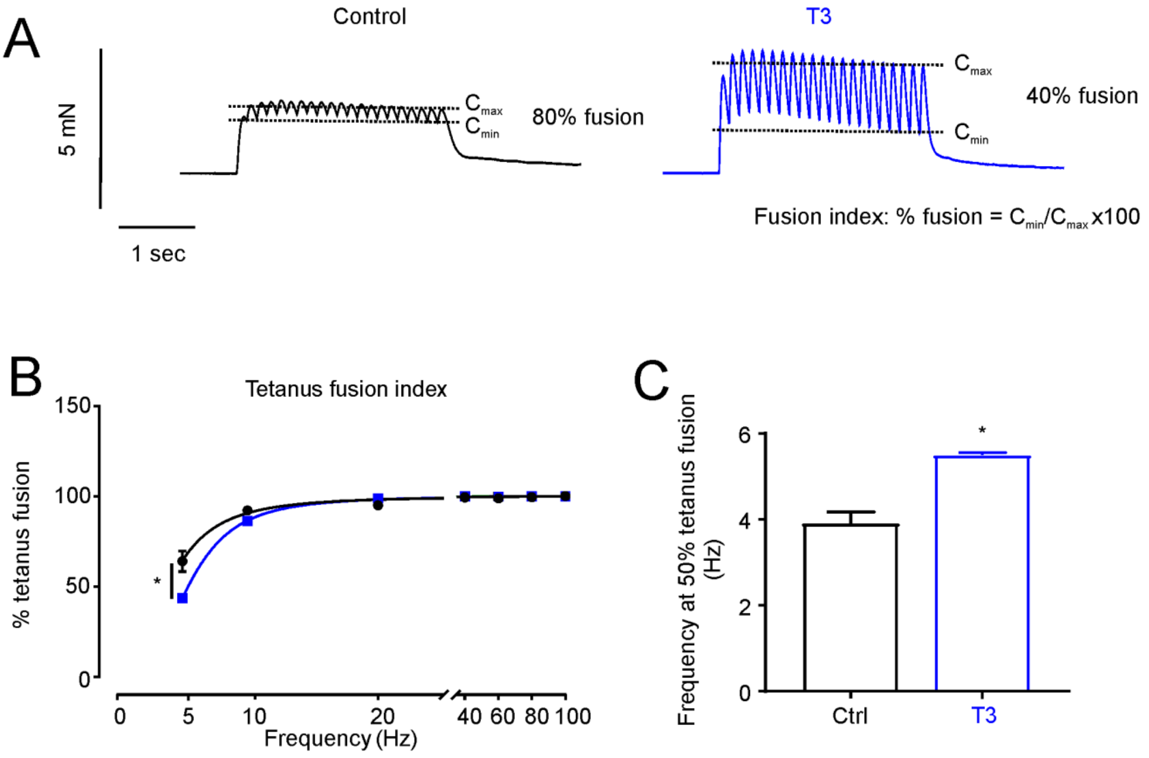


**Supplementary Figure 4 related to Figure 6: Thyroid hormone elevates the tetanus threshold of ESM.**

**A)** Fusion index calculated on representative traces of twitch tension generated by control (black) and +T3 (blue) ESM at 5 Hz tetanus stimulation. The fusion index calculated as the percentage ratio of the maximal relaxation amplitude before the last contraction of the tetanus ($C_{min}$) to the amplitude of this last contraction ($C_{max}$). **B)** The fusion index-frequency curve of control (black line) and +T3 (blue line) ESM. *$p<0.05$ by 2 way-ANOVA and Tukey's multiple comparison test. **C)** Stimulation frequency at 50% tetanus fusion of control (black bar) and +T3 (blue bar) ESM; n = 8/group, *$p<0.05$ by Student's t-test.

### Supplementary Figure 5

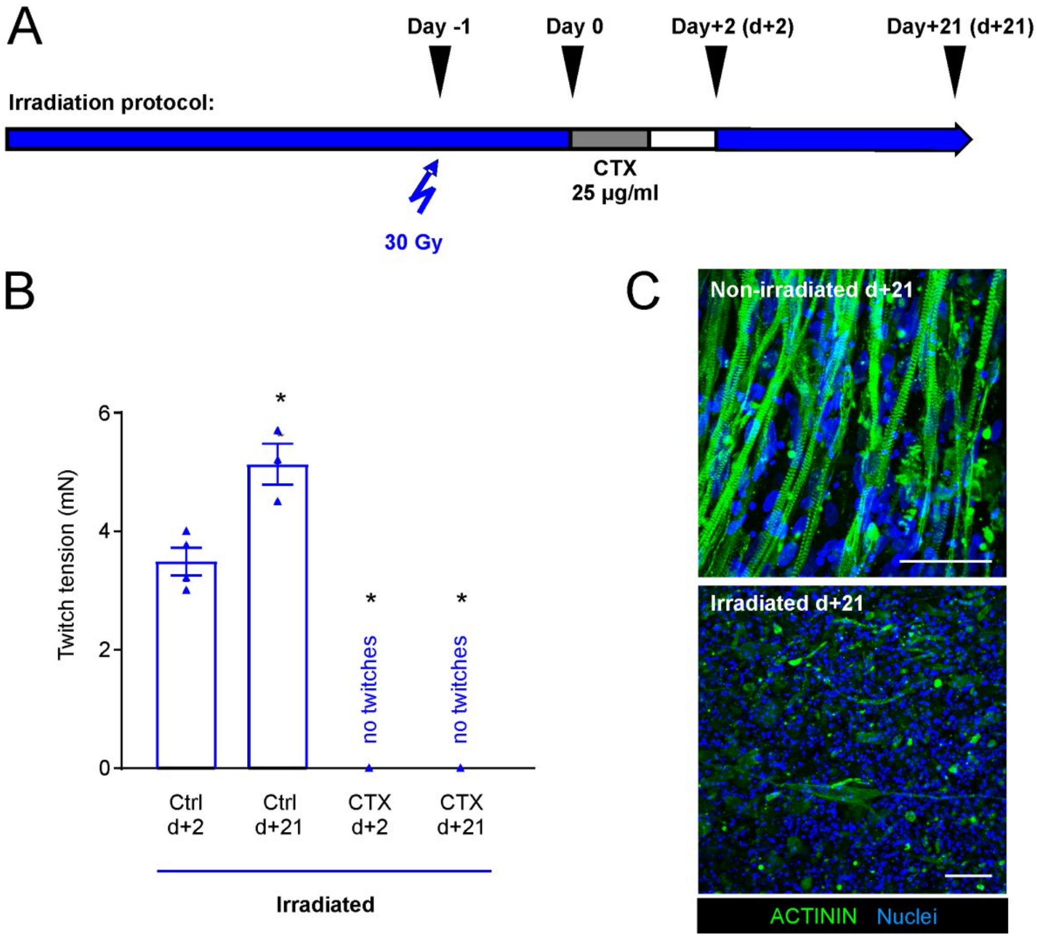


**Supplementary Figure 5 related to Figure 7. Irradiation blocks regenerative capacity of human engineered skeletal muscle. (A)** Experimental scheme of irradiation protocol. One day before cardiotoxin injury ESM were irradiated with 30 Gy. ESM were then incubated with 25 µg/ml CTX for 24 hrs. **(B)** Tetanic twitch tension at 100 Hz stimulation frequency of ESM with irradiation at indicated time points after CTX (25 µg/ml) injury or control (Ctrl) condition; n=3-4/group, *$p<0.05$ vs. Ctrl day+2 by 1-way ANOVA and Tukey's multiple comparison test. **(C)** Immunostaining of sarcomeric $\alpha-$ACTININ (green) and Nuclei (blue) in non-irradiated ESM **(top panel)** and irradiated ESM **(bottom panel)** 21 days after CTX injury. Scale bars: 50 µm.

**Supplementary videos**

**Video 1: Spontaneous contractions of SMO matured for 3 weeks on metal holders**

**Video 2: Spontaneous contractions of ESM matured for 3 weeks on metal holders**

**Video 3: Spontaneous contractions of ESM matured for 11 weeks on metal holders**
